# Avian influenza virus neuraminidase stalk length and haemagglutinin glycosylation patterns reveal molecularly directed reassortment promoting the emergence of highly pathogenic clade 2.3.4.4b A (H5N1) viruses

**DOI:** 10.1101/2024.05.22.595329

**Authors:** Ranjana Nataraj, Anantika Chandra, Sannula Kesavardhana

**Author notes:** Correspondence: Sannula Kesavardhana, Department of Biochemistry, Division of Biological Sciences, Indian Institute of Science (IISc), Bengaluru, Karnataka 560012, India.

## Abstract

The recently emerged panzootic clade 2.3.4.4b H5N1 influenza viruses show unprecedented extensive spread in wild birds and have been transmitted to several mammals, including humans. The virologic factors that have driven the success of the 2.3.4.4b H5N1 viruses, which has not been achieved by previous H5N1 clades, is unclear. We show that the 2.3.4.4b H5 haemagglutinin (HA) paired exclusively with full length (long stalk) N1 neuraminidase (NA) in birds and mammals, unlike previous clades of H5 viruses, which preferentially paired with N1 proteins with stalk deletions (short stalk). We found that the emergence of a 2.3.4.4b H5 HA with seven glycosylation sites was critical in driving its pairing with long stalk N1. The earlier H5 clades that paired with short stalk N1s showed a pattern of eight or more glycosylation sites. A prior shift in a glycosylation site from position 103 to 171 in the receptor binding domain of H5 HA and the subsequent S173A mutation that removed it triggered the emergence of 2.3.4.4b clade H5N1 viruses. Thus, the evolution of novel variations in the H5 HA and their preference for long stalk N1 pairing led to increased fitness and pathogenicity. These observations led us to establish and validate the hypothesis that the pairing of avian virus HA and NA subtypes is not stochastic but is rather molecularly programmed by HA glycosylation and NA stalk length, modulating fitness and emergence of novel avian influenza viruses.

**GRAPHICAL ABSTRACT:** 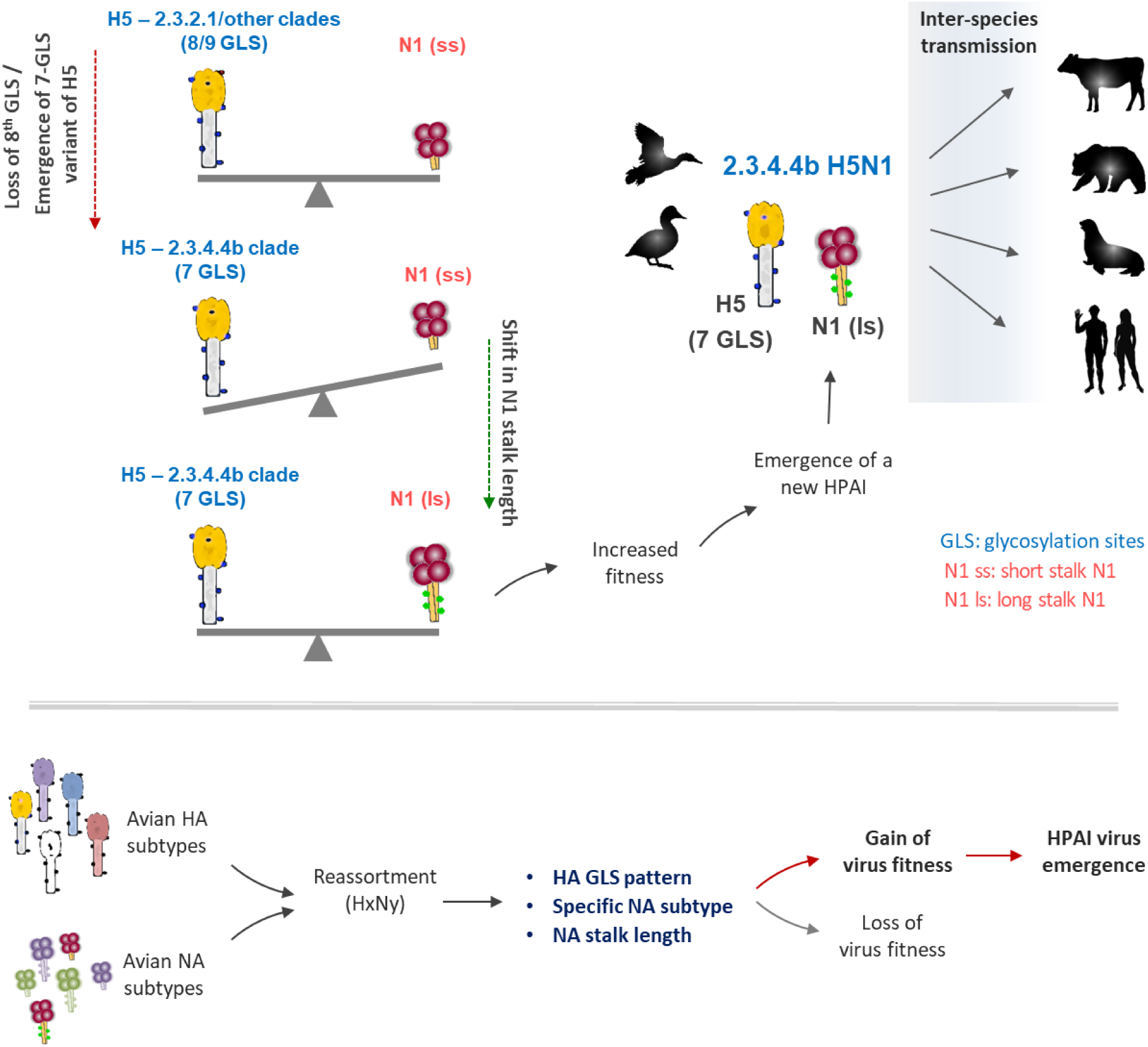

## INTRODUCTION

The influenza A virus (IAV) causes seasonal respiratory infections in humans and occasionally emerges as a pandemic strain. Wild aquatic birds are the primary reservoirs of IAVs and transmit IAVs to diverse species, including humans, swine, seals, horses, and bats (Krammer et al., 2018; Ozawa and Kawaoka, 2013). The subtypes of IAV are classified based on two antigenically essential surface proteins, haemagglutinin (HA) and neuraminidase (NA). To date, 16 different HA and 9 known NA protein variants have been identified in birds, which can reassort and generate diverse avian influenza strains. Most IAVs circulating among wild birds cause mild infections within their natural hosts and are categorized as low-pathogenic avian influenza (LPAI) viruses (Escalera-Zamudio et al., 2020; Yamaji et al., 2020). However, once these viruses cross over to domestic birds, some viruses with H5 or H7 HAs may become more pathogenic, leading to the emergence of highly pathogenic avian influenza (HPAI) viruses (Gilbertson and Subbarao, 2023; Yamaji et al., 2020). Zoonotic transmission of HPAI viruses to humans can cause severe respiratory disease and mortality, although the human-to-human transmission rate is very low (Gilbertson and Subbarao, 2023; Krammer et al., 2018; Ozawa and Kawaoka, 2013). However, the possibility remains that new variations in HPAI viruses may confer adaptation for sustained human-to-human transmission, potentially giving rise to pandemics.

H5Nx (x can be any NA subtype) viruses are predominant HPAI subtypes that circulate between aquatic birds and poultry (Gilbertson and Subbarao, 2023; Wille and Barr, 2022). The H5Nx HPAI viruses frequently reassort with LPAI viruses, leading to the emergence of novel viruses causing seasonal outbreaks in poultry (Escalera-Zamudio et al., 2020; Xie et al., 2023). The current HAs of HPAI H5N1 viruses originated from the goose/Guangdong lineage (gs/GD) that was first reported in 1996 and has been associated with HPAI outbreaks in poultry and human infections (Chen et al., 2006; Subbarao et al., 1998). Several H5 clades have evolved from the gs/GD lineage (Group, 2012; Wille and Barr, 2022), with early clades having a preference for pairing primarily with N1 and N2 NAs. The 2.3.4.4 clade of H5 that emerged after 2011 acquired the ability to pair with several NA subtypes (including N1, N2, N6 and N8) and further diverged into the currently circulating 2.3.4.4b clade after 2014 (de Vries et al., 2015; Global Consortium for and Related Influenza, 2016). For a period of time, the 2.3.4.4b viruses were often H5N6 or H5N8, which was different from most previous H5 clades (Lewis et al., 2021; Li et al., 2021; Wille and Barr, 2022). More recently, the H5N8 viruses which were predominate in many regions of the world, reassorted, leading to the resurgence of the highly pathogenic H5N1 subtype (European Food Safety et al., 2022; Lewis et al., 2021; Xie et al., 2023; Zeng et al., 2024).

The 2.3.4.4b H5N1 viruses have been very successful and have caused persistent outbreaks in wild and domestic birds, leading to millions of deaths worldwide, including in the Americas (European Food Safety et al., 2022; Kandeil et al., 2023; Lewis et al., 2021). The 2.3.4.4b H5N1 viruses have also been detected in an unprecedented range of mammalian species that include farmed minks, sea lions, red foxes, bobcats, black bears, dolphins, humans, and cows (Aguero et al., 2023; Gilbertson and Subbarao, 2023; Maemura et al., 2023; Plaza et al., 2024; Puryear et al., 2023).

It remains unknown why the most recent H5N1 viruses have led to unprecedented persistent outbreaks and spread to diverse species. In this study, we investigated why the 2.3.4.4b H5 paired with N1 and replaced other subtypes across most of the world. We identified that the 2.3.4.4b H5 preferentially paired with glycosylation site-rich full-length (long stalk) N1, unlike previous H5N1 viruses, which paired primarily with N1s with stalk deletions (short stalk). We also establish an intriguing link between the H5 glycosylation pattern and N1 subtype stalk length that might underlie the emergence of the recent 2.3.4.4b clade H5N1 virus.

## RESULTS

### H5N1 gains unusual long stalk N1 in the current circulating 2.3.4.4b clade

To understand the NA preference of avian influenza viruses, we examined the temporal dynamics of various IAV subtypes and their preference for the NA subtype. We observed that N1 and N2 subtypes accounted for nearly 60 percent of the sequenced avian influenza viruses and showed reassortment with H5 or H9 HA subtypes (**Fig. S1A-C**). Also, the circulating IAV strains in swine and humans consisted of mainly N1 and N2 subtypes of NA (**Fig.S1D-E**). Analysis of gs/GD lineage avian influenza viruses showed that the NA subtype varied across the decades and preferentially alternated between the N2 (1990-99) and N1 (2000-09) subtypes of NA (**Fig.1A**). Interestingly, multiple avian NA subtypes (N1, N2, N6, and N8) appeared in the circulation in the following decade (2010-2019), indicating the emergence of NA subtypes that were not in circulation which paired with gs/GD lineage viruses. However, N1 reemerged as a dominant NA subtype in the recent decade by pairing with gs/GD lineage 2.3.4.4b clade H5 HA (**Fig.1A**). This suggests the fitness advantage for the recent 2.3.4.4b clade H5 subtype when paired with N1 subtype despite its ability to pair with other NA subtypes (N2, N6, and N8).

**Figure 1:**
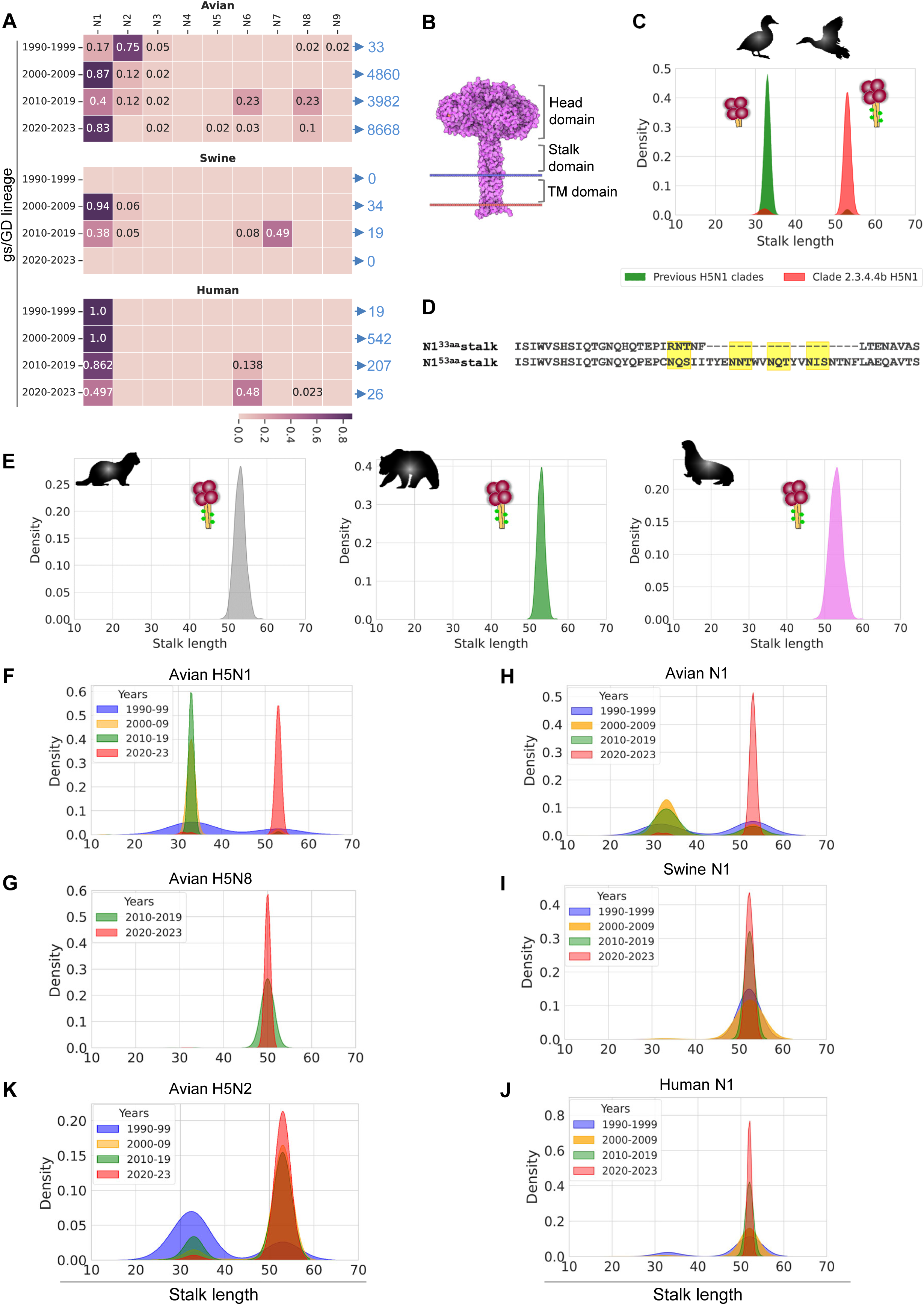
The long stalk N1 prevalence in 2.3.4.4b clade H5N1 infections in birds and mammals. **A**. Decade-wise trends in circulating gs/GD lineage H5 influenza viruses in avian, swine and human hosts represented as heatmaps. Heatmaps are provided for four distinct time periods: 1990-1999, 2000-2009, 2010-2019, and 2020-2023. The number of sequences obtained from GISAID for this decade-wise analysis are indicated in blue. **B.** A model displaying the complete structure of the NA protein and its domain annotation. **C.** Kernel Density Estimate (KDE) plots representing the stalk length distribution of gs/GD lineage avian NA subtype N1 paired with 2.3.4.4b clade H5 HA (2018-2023) or the H5 clades preceding 2.3.4.4b emergence (1990-2017). The x-axis represents the number of amino acids in the N1 stalk domain. The cartoons inside the graph represent short and long stalk versions N1. **D.** Consensus protein sequence alignment of short and long stalks of N1 subtypes in avians. Putative glycosylation motifs (Nx[S/T]) are highlighted in yellow. **E.** KDE plot representing the distribution of N1 stalk lengths in clade 2.3.4.4b H5N1 infections that affected minks, bears, and seals constructed similar to Panel-C. **F, G.** KDE plots illustrating the stalk length distribution for gs/GD lineage avian H5N1 (**F**) and avian H5N8 (**G**). Stalk length distribution from each time period is depicted as a separate curve. **H, I, J.** KDE plots illustrating the stalk length distribution for all avian (**H**), swine (**I**) and human (**J**) N1 subtype of NA from GISAID. **K.** KDE plots illustrating the stalk length distribution for gs/GD lineage avian H5N2.

NA protein consists of a transmembrane (TM) domain, a head domain with sialidase activity, and a stalk domain connecting TM and the head domain (**Fig.1B**). The stalk domain is the most variable region in the NA structure that tolerates insertions and deletions, which determine the viral fitness and adaptation (Blumenkrantz et al., 2013; Castrucci and Kawaoka, 1993; Matsuoka et al., 2009). Notably, NA stalk length influences the transmissibility and host range of IAVs and determines the NA subtype-specific pathogenicity in avian species (Blumenkrantz et al., 2013; Castrucci and Kawaoka, 1993; Matsuoka et al., 2009). We then asked whether NA stalk length plays a role in the emergence of 2.3.4.4b H5Nx strains and determine the fitness of circulating avian strains. Interestingly, all the gs/GD lineage H5N1 viruses that appeared before the emergence of 2.3.4.4b clade (before 2020) were preferably associated with N1 with a short stalk of 33 amino acids (**Fig.1C-D**). However, the current circulating 2.3.4.4b clade H5 showed strict pairing with the N1 subtype with a long stalk of 53 amino acids, as evidenced by the density function (**Fig.1C-D**). The N1 with the long stalk showed an additional 20-residue stretch constituting four putative glycosylation motifs (Nx[S/T]) (**Fig.1D**). Recent reports show that 2.3.4.4b clade H5N1 has spread across all the continents and successfully transmitted to diverse mammalian species (Aguero et al., 2023; Burrough et al., 2024; Plaza et al., 2024; Puryear et al., 2023; Uyeki et al., 2024). We analysed the N1 stalk lengths in mammals infected exclusively with 2.3.4.4b clade. We found that minks, black bears, sea lions, bobcats, red foxes, and dolphins were infected with the 2.3.4.4b clade H5N1 with the long stalk, and no infections were showing short stalk N1 (**Fig.1E & Fig.S2A**). The recent penguin infections also showed long stalk N1. In addition, the analysis of recent H5N1 infections in humans and cows showed that they were infected with 2.3.4.4b clade H5N1 with long stalk N1 (**Fig.S2B**). Thus, 2.3.4.4b clade H5N1 with long stalk is a highly adaptable strain across species.

The decade-wise gs/GD lineage avian influenza sequences revealed that the N1 stalk length distribution was notably bimodal across all decades, with two distinct peaks, representing short and long stalks (**Fig.1F**). The circulating gs/GD avian influenza N1 subtype predominantly consisted of a short stalk until 2019 (**Fig.1F**). Thus, contrary to the N1 subtype of previous gs/GD lineage clades that preferred short stalk length, the N1 of 2.3.4.4b clade H5N1 strain that recently emerged preferred a long stalk length. Also, the N8 subtype consistently showed the long stalk in the recently emerged 2.3.4.4b clade H5N8 (**Fig. 1G**). These observations indicate that the currently circulating 2.3.4.4b clade H5 HA prefers to reassort with the N1 or N8 with a long stalk to confer fitness and emergence. Analysis of all the avian influenza sequences from GISAID indicated that the N1 subtype consisted of a short stalk until 2019, regardless of the HA type it paired, similar to gs/GD lineage pattern (**Fig.1H**). Unlike avian influenza, human and swine IAVs preferred long stalk length N1 across the decades (**Fig.1I-J**). This discrepancy in stalk length of the N1 subtype of NA might be one of the contributing factors in limiting spillover events of IAVs. However, the ability of the long stalk N1 variant to pair with the recent gs/GD lineage 2.3.4.4b clade H5 resulted in the convergence of N1 stalk length preference across hosts, which could facilitate the zoonotic transmission and productive infections of 2.3.4.4b clade H5N1.

In contrast to N1, the gs/GD lineage avian N2 subtypes consistently exhibited a longer stalk length (**Fig.1K**). Notably, only 16 infections (2018-2023) were reported having the N2 subtype that paired with 2.3.4.4b clade H5 in GISAID, indicating its inability to establish itself as a dominant circulating strain. Analysis of all the avian influenza sequences with N2 (irrespective of the HA type) showed two variants of long stalk distinguished by a putative glycosylation site (**Fig.S2C-D**). Also, acquiring this glycosylation site is correlated with a reduction in N2 subtype influenza viruses in circulation (**Fig.S2C-D**). Overall, these observations indicate that the NA stalk length trend in gs/GD lineage avian N1 and N2 subtypes was mutually exclusive until the appearance of long-stalked N1 in the 2.3.4.4b clade H5N1. The recent shift in the distribution of stalk lengths observed in the avian N1 subtype indicates the functional significance of NA stalk length in the emergence of new avian influenza strains.

In addition to the long stalk, we found that the N1 paired with the 2.3.4.4b clade H5 HA showed several mutations in the head domain compared to the previous N1 subtype (**Fig.S3A-D**). None of these mutations span to the sialic acid binding site in N1 (**Fig.S3D-F**). We modeled the structure of the head domain with all the variations present in the recent N1 and found that these mutations did not perturb the overall topology of the head domain and the sialic acid binding site conformation (**Fig.S3A-C**). A few substitutions, V256M, S257P, and A312E, were in proximity to the sialic acid binding residues and appeared not to affect the sialic acid binding conformation (**Fig.S3E-F**). These observations further suggest the critical role of the stalk domain of N1 in the emergence of 2.3.4.4b clade H5N1.

### N1 displaces N8 as a preferred partner of 2.3.4.4b clade H5 and shows increased fitness

Once clade 2.3.4.4b H5 appeared in 2014, it reassorted with the N8 NA with a long stalk and emerged as pathogenic H5N8 (Lewis et al., 2021; Xie et al., 2023). This marked a deviation from the previous clade H5 (2.3.2.1), which predominantly paired with N1 with a short stalk (**Fig.S4A-B**). In 2020, clade 2.3.4.4b H5N8 underwent reassortment with low pathogenic avian influenza (LPAI), resulting in the emergence of H5N1, H5N6, and various other strains (Gilbertson and Subbarao, 2023; Xie et al., 2023). H5N1 emerged as the predominant strain, constituting nearly 81% of reported clade 2.3.4.4b H5 cases in 2020 (**Fig.S4C**). What factors contributed to the increased fitness of clade 2.3.4.4b H5N1 over H5N8 during 2020-2023? In contrast to the N1 subtype, N6 and N8 stalk lengths remain unchanged (**Fig.S4B & S4D**). In the 2014-2019 period, N8 and N6 were paired with 2.3.4.4b clade H5; however, the N8 or N6 pairing could not become a dominant subtype when the analysis was extended to all the H5 clades (**Fig.S4A**). Interestingly, when the 2.3.4.4b clade H5 shifted from N8 to N1 pairing, the resulting 2.3.4.4b H5N1 subtype was dominant irrespective of the H5 clade (**Fig.S4C**). Also, there was a sudden shift in N1 stalk length when it paired with 2.3.4.4b, unlike N6 and N8 (**Fig.S4D**). Perhaps the 2.3.4.4b clade-specific mutations in H5 provided a fitness advantage when paired with N1 compared to N6 and N8.

### Altered glycosylation site pattern and the emergence of seven glycosylation sites variant of H5 HA is linked to the pairing of 2.3.4.4b H5 with long stalk N1

The gain of the long stalk in N1 correlated with the emergence of currently circulating 2.3.4.4b clade H5N1. Therefore, the current circulating avian H5N1 likely exhibits altered neuraminidase activity due to the acquisition of long stalk N1. The HA and NA coevolution is known to maintain the functional equilibrium of sialic acid affinity and sialidase activity essential for virus entry and exit (Kosik and Yewdell, 2019; Liu et al., 2022; Wagner et al., 2002; Xu et al., 2012). Also, the functional balance of HA and NA is critical for conferring virus fitness and virulence. Thus, if the circulating N1 demonstrates altered neuraminidase activity, the accompanying H5 must possess compensatory adaptations. Our comparison of the amino acid sequences of currently circulating avian H5 HA (2018-2023), paired with short stalk and long stalk N1, revealed specific variations confined to the receptor binding site and its surrounding regions (**Fig.2A & Fig.S5A**). A clade-wise distinction of H5 indicated that short stalk N1, currently in circulation, preferentially paired with previous H5 clades (2.3.2.1a and 2.3.2.1c), with a small fraction of short stalk N1 pairing with the current 2.3.4.4b clade (**Fig.2B & Fig.S5B**). On the other hand, long stalk N1 exclusively paired with the 2.3.4.4b H5 clade but not with previous H5 clades (**Fig.2B**). Further analysis of 2.3.4.4b clade H5 HA that paired with short and long stalk N1 subtype unveiled two specific variations in the receptor binding domain (RBD) at residues 173 (S➔A) and 186 (Q➔R)) (**Fig.2C**). These variations were also conserved (S173 and Q186) in previously dominant H5 clades (2.3.2.1a and 2.3.2.1c) that paired with short stalk N1 when compared to the long stalk N1. In the H5 HA structure, the S173 residue spans the loop region, which is a part of the RBD, and Q186 is distantly located from the RBD (**Fig.2C**). The S173A substitution in H5 HA that paired with long stalk N1 disrupts a glycosylation motif, which might impact RBD accessibility for binding to sialic acids (**Fig.2C**). This revealed the previously unknown specific glycosylation motif variation in 2.3.4.4b H5 HA that paired with short and long stalk N1. The S173A substitution in RBD prompted us to investigate how the H5 HA glycosylation pattern varies in pairing with short and long stalk N1. Strikingly, we found distinct glycosylation site patterns in the H5 HA that paired with short or long stalk N1 (**Fig.2D**). The 2.3.4.4b clade H5 HA that paired with long stalk N1 (most prevalent) showed preferably 7 glycosylation sites, however, it had 8 or 9 glycosylation sites when paired with short stalk N1 (less prevalent) (**Fig.2D-E**). Also, H5 HA showed 8 or 9 glycosylation site patterns when paired with short stalk N1 independent of the clade. Thus, these observations indicate an intriguing correlation between H5 HA glycosylation site distribution and the stalk length of the N1 in circulating avian influenza strains.

**Figure 2:**
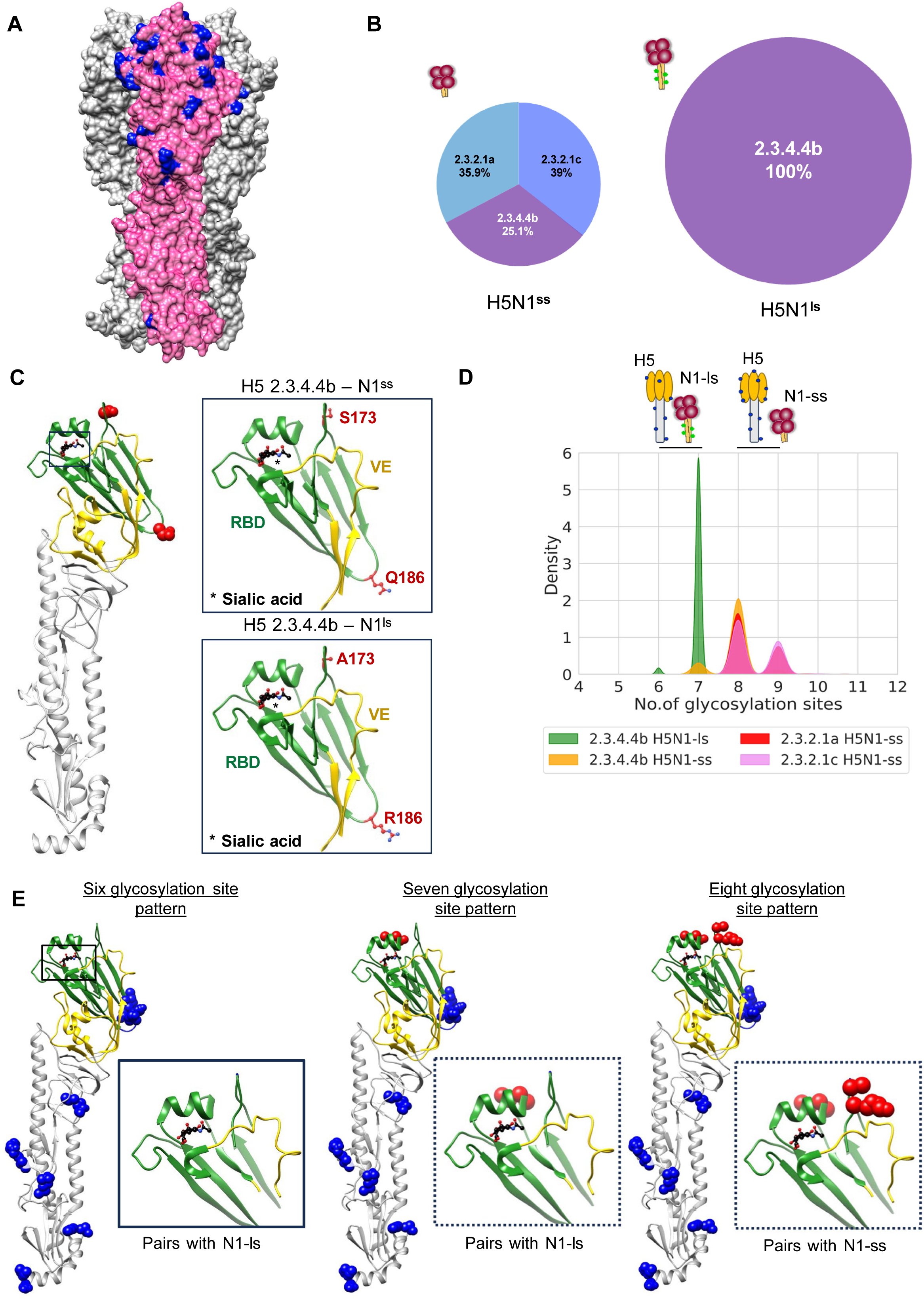
The glycosylation site patterns of avian H5 HA molecularly linked to its pairing with specific NA subtype. **A.** Surface representation of H5 trimer (PDB: 3UBE), with a monomer structure highlighted in magenta. The surfaces of the other two monomers are shown in gray. The residues highlighted in blue indicate the positions that vary between H5 pairing with short stalk and long stalk N1 subtype present in 2018-2023. **B.** Pie charts depicting the H5 clade distribution within H5N1 sequence deposited in GISAID from 2018 to 2023. The charts are differentiated based on the stalk length of the N1 subtype of NA (on left H5N1 with short stalked N1; on right H5N1 with long stalked N1). Distinct clades are color-coded, and annotations display the percentage of sequences in each clade. The size of each pie chart represents the abundance of sequences with the corresponding N1 stalk length feature. **C.** Monomeric H5 structure highlighting the two amino acid positions differing in 2.3.4.4b clade H5 HA that pairs with a short stalk or long stalk N1 represented as spheres. The Receptor Binding Domain (RBD) and Vestigial Esterase (VE) domain are colored green and yellow, respectively. Insets depict a zoomed-in view of the RBD domain and the amino acids at these two positions. **D.** KDE plot illustrating the number of putative glycosylation sites in H5 HA of clades 2.3.2.1a, 2.3.2.1c, and 2.3.4.4b paired with short stalk (ss) vs. long stalk (ls) N1 subtype. **E.** Annotation of glycosylation site variants (6th, 7th, and 8th) of H5 HA from Panel-D, shown on the monomeric H5 structure (PDB: 3UBE). Glycosylation positions are represented as blue spheres, with the RBD and VE domains highlighted in green and yellow, respectively. The 7th and 8th glycosylation sites are shown in red.

We further analysed the glycosylation motif profile of avian influenza H5 HA from 1990 to 2023 to investigate whether H5 HA glycosylation and NA stalk length correlation occurred across decades. Avian H5 strains paired with short stalk N1 revealed three distinct glycosylation site patterns (**Fig.3A**). The most common pattern comprised 7 or 8 glycosylation sites (hereafter referred as 7-GLS and 8-GLS) in H5 HA, with some variants showing 9 glycosylation sites. In contrast, H5 paired with long stalk N1 exhibited only two glycosylation patterns, with the prevalent variant retaining 7-GLS and a small population featuring 6 glycosylation sites. Notably, the H5 subtype with 8-GLS did not pair with long-stalked N1 and preferred only short-stalked N1 (**Fig.3A**). This suggests the molecular link between H5 HA with 7-GLS (deglycosylated variant) and long stalk N1 that confer reemergence and fitness of 2.3.4.4b clade H5N1 strain.

**Figure 3:**
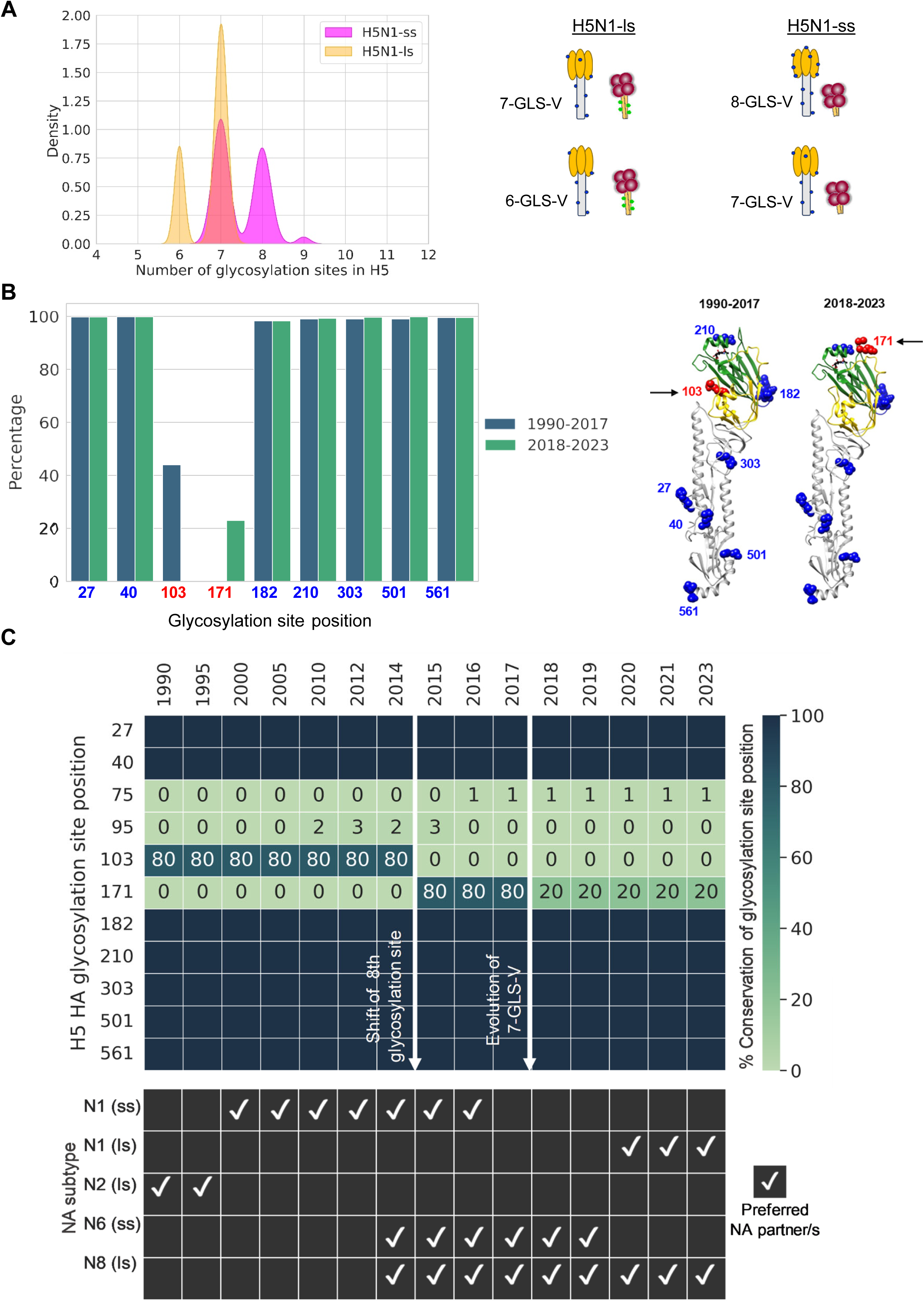
A shift in 8^th^ glycosylation site in RBD of H5 HA and the evolution of seven glycosylation site variant of H5 are a prerequisite for gaining long stalk N1 in 2.3.4.4b clade. **A.** KDE plot illustrating the number of putative glycosylation sites in H5 HA paired with short stalk or long stalk N1 subtype from 1990 to 2017. ls-long stalk; ss-short stalk; GLS-V – glycosylation site variant. **B.** The degree of conservation of the glycosylation sites of H5 HA for the time periods of 1990-2017 and 2018-2023, represented as bar plots. The glycosylation site positions (in H3 numbering) are on the x-axis. The 8^th^ glycosylation site variants of H5 and their position in RBD are shown in ribbon structure (PDB:3UBE). The 7 conserved glycosylation sites are shown as blue spheres and the 8^th^ glycosylation site which differs between the two datasets are labeled in red. The RBD and VE domains of the monomeric structure are highlighted in green and yellow, respectively. **C**. Heatmap illustrating the percent conservation of each glycosylation site of the H5 HA obtained from a specific time frame as indicated in the plot. The horizontal axis denotes the years sampled, and the vertical axis represents the glycan positions of H5 in H3 numbering. The bottom panel displays the prevalent NA subtype and its stalk length feature that H5 paired with during the respective year.

The discrepancies in the H5 HA glycosylation pattern primarily stem from the Vestigial Esterase (VE) domain (residues 103-105; represents the 8^th^ glycosylation site) and the RBD (residues 210-212, represents the 7^th^ glycosylation site) of the HA head domain (**Fig.3B&2E**). Mapping these glycosylation patterns onto the H5 HA structure revealed that the positioning of seven of these sites is conserved across the clades. However, the 8^th^ glycosylation site appeared to be variable that shift between 103 and 171 glycan positions (**Fig.3B, left panel**). Earlier clades of H5 HA harbored the 8^th^ glycosylation site in the VE domain (residues 103-105), but this site was shifted to the RBD in the current circulating 2.3.4.4b clade H5 (residues 171-173) (**Fig.3B**). Despite the presence of 8-GLS variant of H5 HA, the most prevalent H5 HA in this decade showed only 7-GLS pattern, which paired with long stalk N1. 8-GLS variant of H5 HA preferentially pairs with short stalk N1 conferring functional balance and viral fitness of H5N1 in previous clades. Perhaps the newly emerged 8-GLS variant of H5 HA, which showed a shift of the 8^th^ glycosylation site to RBD, might have posed a fitness cost when paired with short stalk N1 and thus might destabilize the functional balance of HA and NA.

To understand the evolution of the novel 7-GLS variant of 2.3.4.4b clade H5, we examined avian influenza H5 HA glycosylation site profiles. We found that the shift in 8^th^ glycosylation site in the RBD of H5 HA occurred during 2014 (**Fig.3C**). However, the new 8-GLS variant of H5 dominated the circulation of the avian influenza strains for a transient period and preferentially paired with short stalk N1 or N6 and N8 (**Fig.3C**). In 2017, the H5 variant of 7-GLS showing S173A mutation emerged, which dominated over the new H5 variant of 8-GLS and paired strictly with long stalk N1 (**Fig.3C**). Thus, a shift in the 8^th^ glycosylation site (103-105 ➔ 171-173) followed by the emergence of the variant with 7-GLS preceded the acquisition of long-stalked N1 likely to retain HA-NA functional balance. The pairing of 7-GLS variant of 2.3.4.4b H5 with the N1 with shorter stalk length could disrupt the functional balance of H5 and N1, resulting in decreased viral fitness. We further extended the analysis of glycosylation sites in H5 clade influenza viruses in domestic and wild aquatic birds to understand the specific avian hosts facilitating the shift in 8^th^ glycosylation site and the emergence of 7-GLS variant of 2.3.4.4b clade H5. It appeared that the 8th glycosylation site shift (103-105 ➔ 171-173) in the RBD of H5 HA and the subsequent emergence of 2.3.4.4 H5 with 7-GLS occurred in domestic birds (**Fig.S6A**). The glycosylation motif positions 103 and 171, representing the 8th glycosylation site in H5 HA, were conserved in domestic birds, but these positions were not conserved in wild aquatic birds (**Fig.S6A-B**). Instead, residue positions 75 and 95 represented 8^th^ glycosylation site in wild aquatic birds, although these glycosylation sites were not prevalent (**Fig.S6B**). Thus, the shift in 8^th^ glycosylation site and the emergence of 7-GLS variant of H5 HA were confined to domestic birds (**Fig.S6A-B**). Recent studies indicate that the 2.3.4.4b clade H5N1 acquired the NA from wild birds (Jimenez-Bluhm et al., 2023; Kandeil et al., 2023; Wille and Barr, 2022; Xie et al., 2023). Our observations indicate that the transmission of long stalk N1 of LPAIs from wild birds to domestic birds might have led to the pairing of H5 7-GLS (from domestic birds) with long stalk N1 (originated from wild birds).

To further understand the correlation of the H5 glycosylation site pattern and its pairing with long stalk N1, we checked the glycosylation pattern of H5 in H5N8 from which the recent H5N1 originated. We found that H5 of 2.3.4.4b clade paired with N8 (H5N8) in 2018 showed a loss of glycosylation site and showed only 6 or 7 glycosylation patterns (**Fig.S7A-B**). Also, N8, paired with H5, showed a long stalk length. However, the H5 paired with N6 showed 7 and 8-GLS patterns. This further strengthens our hypothesis that the possible deglycosylation-driven emergence of 7-GLS variant of 2.3.4.4b clade H5 could be the driving force for pairing with long stalk NA.

In H5 variants of 6 and 7 glycosylation sites, the T105E mutation was prevalent, which disrupts 8^th^ glycosylation site. The glutamic acid residue at position 105 (E105) likely forms hydrogen bonds with asparagine at 73 position (N73) of the HA stalk, stabilizing the adjacent HA stalk (**Fig. S7C**). The loss of the 8^th^ glycosylation site (residues 103-105) may have contributed to enhanced HA trimer stability in H5 HA paired with long stalk N1. Additionally, in the 6 glycosylation sites variant, the glycosylation site in the RBD corresponding to the 7^th^ glycosylation site is lost (210-212), resulting in the unobstructed exposure of the sialic acids binding pocket, potentially facilitating enhanced accessibility for sialic acid binding (**Fig. 2E & 3B**). The observed loss of 8^th^ glycosylation site in the VE domain and less frequent loss of 7^th^ glycan site in the RBD of H5 might enhance the sialic acid binding affinity of H5. Thus, these observations indicate an evolutionary link between the stalk length distribution of N1 and the glycosylation site coverage of H5, which confers fitness advantage and the emergence of new H5Nx variants.

### H5 glycan surface plasticity governs its reassortment with specific NA subtype

Our analysis established a correlation between H5 HA glycosylation pattern and its pairing with N1 subtype having long or short stalk length. A time series analysis of H5Nx sequences showed that at any given time, H5 pairs with multiple NA subtypes and the preferred NA partner was not consistent (**Fig.4A**). In certain decades N1 was the most common partner while in other decades, N2 and other NA subtypes emerged as the preferred partner (**Fig.4A**). Given that HA-NA pairing is successful only when the sialidase activity of NA and sialic acid affinity of HA is functionally balanced, avian H5 HA must be able to modulate sialic acid affinity, allowing it to pair with multiple NA types. Modifying glycosylation coverage is one of the critical strategies adopted by HA to modulate its sialic acid affinity (de Vries et al., 2010; Wang et al., 2009). Thus, we hypothesized that multiple H5 HA subpopulations with distinct glycosylation site patterns could exist at any given timeline, and they could be sampled by different NA types. The glycosylation coverage most common in the H5 population in the given season/time might possibly determine the prevalent NA subtype with which H5 HA pairs.

**Figure 4:**
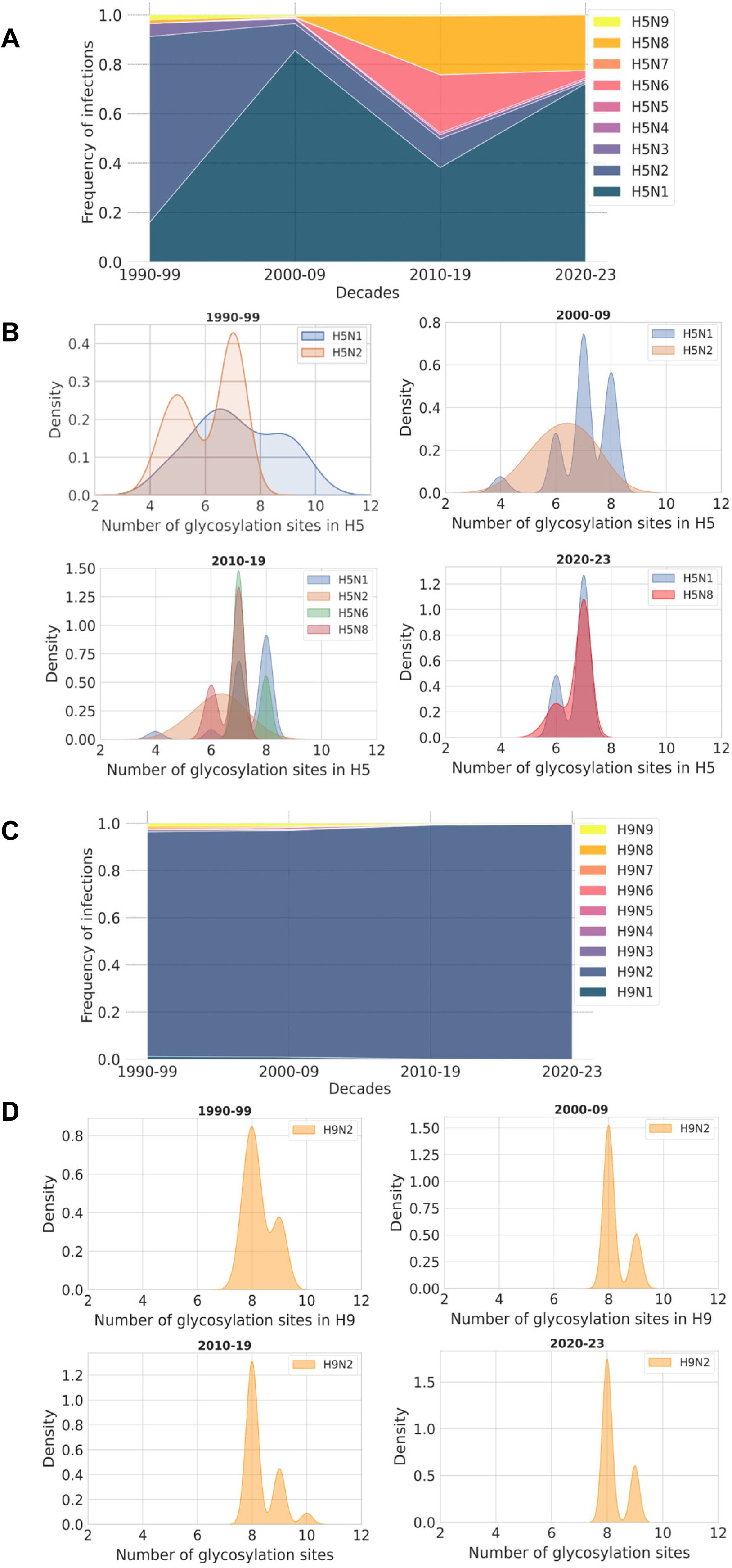
H5 HA glycan surface plasticity determines its reassortment with specific NA subtype. **A.** Cumulative time series plot illustrating the evolution of NA subtype preference of H5 HA. The y-axis represents the frequency of H5Nx cases, with each trace colored based on the NA subtype. **B**. KDE plots supplementing the time series plots in Panel-A, displaying the glycosylation site distribution of H5 HA in each decade. The curves in the density plot are color-coded according to the NA subtype with which H5 pairs. **C.** Cumulative time series plots illustrating the NA subtype preferences of H9 HA, constructed similar to Panel-A. **D**. KDE plots for Panel-C, displaying the glycosylation site distribution of H9 HA in each decade.

During 1990-99, avian H5 was predominantly associated with the N1 and N2 subtype of NA, with N2 pairing being more common (**Fig.4A**). An analysis of glycosylation sites in H5 during this period revealed that the variant of 7-GLS was most prevalent, followed by the 5, with smaller fractions exhibiting 6 and 8 glycosylation sites (**Fig.4B**). Examining the corresponding NAs, we noted that the N2 subtype preferred to pair with 7 and 5 glycosylation sites variant of H5, while N1 paired with 8 and 7-GLS variants (**Fig.4B**). The prevalence of variants of 5 and 7 glycosylation sites, effectively balancing NA activity of N2 subtype, likely contributed to N2 subtype being the preferred H5 partner in that decade and the dominance of H5N2 in circulating strains. In the subsequent decade of 2000-09, H5 paired with N1 and N2 subtypes of NA, with N1 becoming the preferred partner of H5 HA. Notably, the H5 variant of 5 glycosylation sites that paired with N2 showed a decline (**Fig.4B**). Interestingly, the H5 variants of 7 and 8-GLS, preferred by the N1 subtype, became more common. The decrease in the H5 variant of 5 glycosylation sites that paired with N2, coupled with the increased prevalence of the variants of 7 and 8-GLS, might underlie the shift towards N1 as the preferred partner in that decade. The N1 subtype that became prevalent in this decade, which paired with H5 variants of 7 and 8 -GLS, showed short stalk length. In the decade of 2010-19, which marks the emergence of the 2.3.4.4 clade, H5 expanded its ability to sample multiple NA subtypes. The H5 clades in this decade showed pairing with N6 and N8 subtypes along with N1 and N2 (**Fig.4B**). The H5 variant of 8-GLS, circulating in the previous decade, showed a decline, and the 6 and 7-GLS variants became dominant (**Fig.4B**). The decline in H5 variant of 8-GLS in circulation possibly contributed to the decrease in N1 subtype pairing. The loss of the 5 glycosylation sites variant in this decade further supports the decreased pairing of H5 with the N2 subtype. N6 and N8 displayed a preference for the H5 variant of 7-GLS, the most common in that decade. In the current decade (2020-23), the H5 variant of 8 glycosylation sites disappeared, and only the H5 with 6 and 7-GLS variants sustained in circulation (**Fig.4B**). Intriguingly, the N1 subtype exhibited a strong preference for the H5 variants of 6 and 7-GLS, while N8 continued to pair with the 7-GLS variant. This striking shift in N1’s preference to the H5 variants of 6 and 7-GLS H5 instead of 8-GLS was associated with the emergence of long stalk N1, which was not seen in previous clades. Both N1 and N2 subtypes encountered a decline in their preferred H5 partners having 5 or 8-GLS. However, the N1 subtype appeared to have adapted by gaining the long stalk, enabling its preference for the dominant H5 variant with 7 glycosylation sites. In contrast, N2 did not show alteration in its stalk length, potentially explaining why N1 re-emerged as an H5 partner while N2 pairing did not rebound after the decline in H5 variant of 5 glycosylation sites. Thus, contrary to the assumption that the pairing of specific NA subtypes with H5 is random, our data suggest a molecular signature that governs the pairing of H5 HA with specific NA subtypes.

To further corroborate the role of avian HA glycosylation pattern association to the NA subtype pairing, we extended our analysis to the avian LPAI subtype H9 (**Fig.4C-D**). The H9 subtype is the highly prevalent HA type circulating within avian hosts. Interestingly, H9 appeared to exclusively pair with N2 over the decades, in contrast to H5, which exhibited a dynamic interaction with multiple NA types concurrently (**Fig.4C**). Our observations indicated that the glycan surface diversity allowed H5 to interact with various NA subtypes, and the preferred NA partner of H5 varies based on the glycosylation site profile of H5 in circulation. Also, our hypothesis suggests that an avian influenza HA type maintaining a stable glycosylation profile would consistently pair with one NA partner. Limited glycosylation site variants would restrict its ability to interact with multiple NA types. We observed that avian H9 predominantly displayed 8 glycosylation site variants, with only a few instances of 9 and 10 glycosylation sites (**Fig.4D**). Examining the evolution of the glycosylation sites of H9 across time revealed a stable distribution of glycosylation site pattern, with the 8 glycosylation sites variant consistently being the most prevalent. This restricted H9 pairing with N2 subtype across the decades (**Fig.4D**). Overall, these observations unraveled an evolutionary link between the glycosylation pattern plasticity of avian HA with specific NA subtype pairing and the emergence of dominant circulating avian IAV strains.

### The glycosylation pattern of HA and stalk length of NA are the determinants of fitness and emergence of avian influenza subtypes

16 HA (H1-H16) and 9 NA (N1-N9) subtypes currently circulate in avian species and any given HA subtype would be able to reassort with all NA subtypes and vice versa. The pairing of HA and NA was considered a random phenomenon, and the evolution of reassorted strain in a particular host is determined by the HA and NA functional balance of sialic acid receptor binding and receptor destroying activities. Our observations here revealed an intriguing link between the glycosylation pattern of HA and the stalk length of NA, which determines the fitness of avian influenza strains. We argue that specific structural features of avian HA and NA regulate the evolution of dominant circulating strains. Thus, we hypothesize that the reassortment resulting in HA and NA pairing and the emergence of dominant circulating avian influenza strains is not a stochastic event and is governed mainly by the glycosylation profile of HA, the type and stalk length of NA. To substantiate this hypothesis, we constructed a multiparametric bubble plot visualizing the variation in glycosylation sites of H5 HA, the NA subtypes, and the NA stalk length distribution in a single plot using H5Nx sequences across all decades. The multiparametric bubble plot indicated that the pairing of an H5 HA with a particular glycosylation site pattern was restricted to a specific NA subtype with a defined stalk length (**Fig.5A**). A higher number of glycosylation site variants of H5 enabled its pairing with diverse NA subtypes with distinct stalk lengths. Also, H5 with 5 or less glycosylation sites appeared to show a very low pairing frequency with any given NA subtypes.

**Figure 5:**
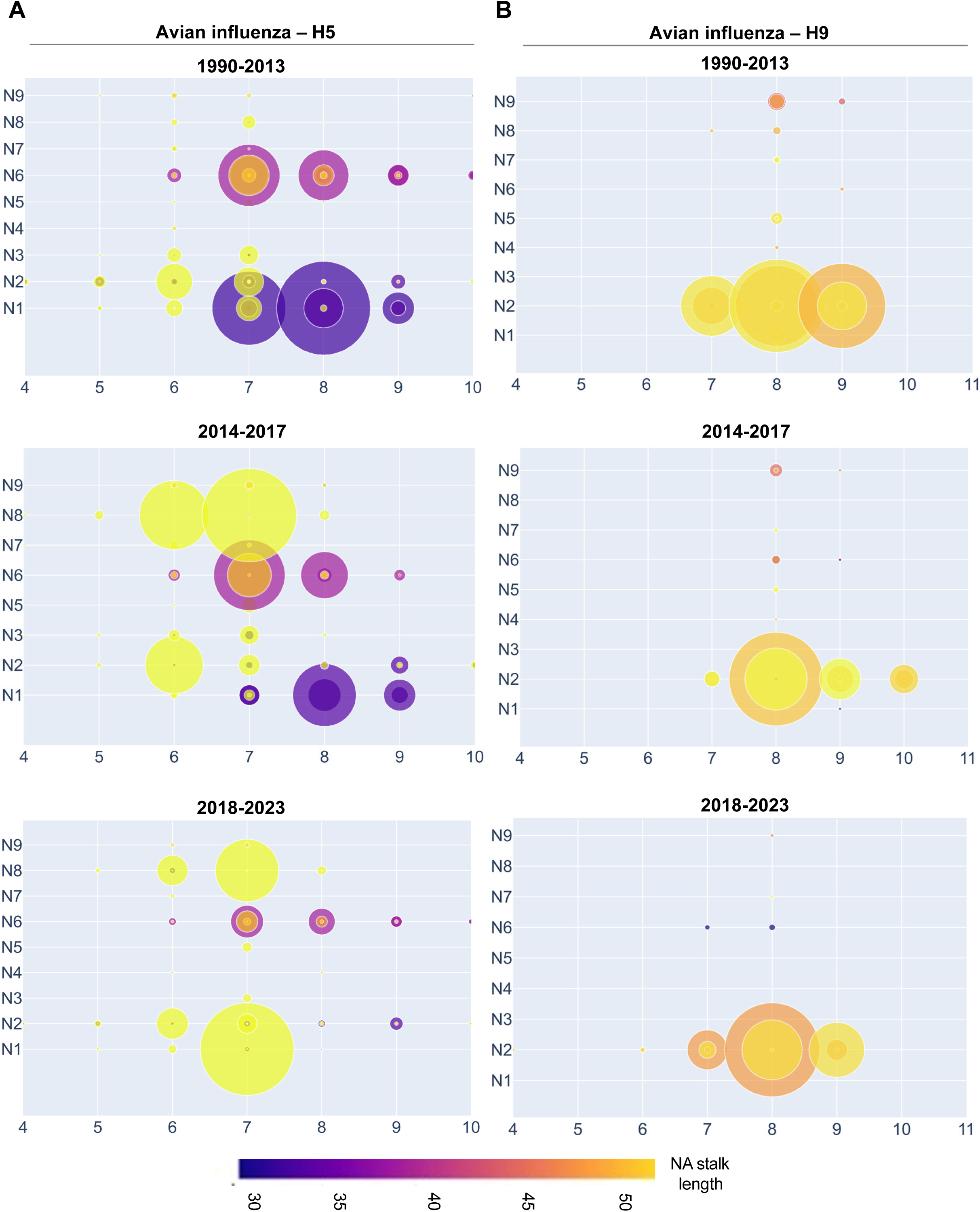
Specific molecular features program the fitness and emergence of avian influenza viruses. **A**. Multiparametric bubble plots illustrating the interplay between H5 glycosylation sites distribution, NA stalk length, and H5Nx pairing at time periods as indicated. Each point in the plot denotes four parameters: the x-axis value denotes the number of glycosylation motifs in H5, the y-axis value denotes the NA subtype, and the size of the bubble is proportional to the frequency of that particular H5Nx combination. The color of the bubble reflects the stalk length of the NA in the H5Nx combination. **B**. Multiparametric bubble plots illustrating the H9 glycosylation sites distribution and its association with specific NA subtype and its stalk length, similar to Panel-A.

To establish the pairing principles that led to the emergence of the 2.3.4.4b H5N1 clade, we constructed bubble plots representing the time scales before and after the emergence of the 2.3.4.4 clade (1990-2013 and 2014-2017) or 2.3.4.4b (2018-2023) subclade (**Fig.5A**). We observed mainly 7 or 8-GLS variants of H5 HA before the emergence of 2.3.4.4 clade H5 that predominantly paired with short-stalk N1 (as evidenced by the large radii of the bubble). These H5 variants were also paired with short-stalk N6 at a moderate frequency. In contrast, variants of 6 or 7 glycosylation sites became more prevalent than H5 having 8 glycosylation sites after the emergence of 2.3.4.4 clade H5 (**Fig.5A**). These H5 variants were reassorted highly with the long stalk N8 subtype and long stalk N2 with moderate frequency. The 8-GLS variant of H5 paired with only short stalk N1 or N6 subtypes (**Fig.5A**). Thus, H5 HA with a higher number of glycosylation sites prefer short stalk N1 or N6 subtypes, and loss of glycosylation sites in H5 shifts preference for long stalk NA subtypes. H5 HA variant with 7-GLS appeared as a highly prevalent subtype during 2.3.4.4b subclade emergence, unlike the previous clades of H5 that show multiple variants of glycosylation sites (**Fig.5A**). The H5 variant with 7-GLS paired preferentially with long stalk N1 and moderately with long stalk N8 subtypes. The emergence and dominance of H5 variants with 7-GLS appeared to have restricted short stalk NA subtypes in circulation, and only long stalk N1 and N8 subtypes dominated circulating strains. Thus, H5 HA with 7 or fewer glycosylation sites prefers to pair with long stalk N1 and N8 than any other NA subtypes. To validate our hypothesis further, we extended our analysis to the highly prevalent H9 subtype in the avian population. Interestingly, the H9 subtypes predominantly had 8 glycosylation sites, and a variant of 9 glycosylation sites also existed but to a lesser extent (**Fig.5B**). Since its glycosylation pattern is restricted over time, the H9 was paired with only the long stalk N2 subtype of NA across all the decades (**Fig.5B**). Our analysis of H5 and H9 reassortment patterns with NA subtypes establishes that monitoring avian HA glycosylation site patterns and specific NA subtypes and their stalk lengths enables the possible prediction of the emergence and fitness of novel avian influenza strains.

## DISCUSSION

Avian influenza viruses cause seasonal outbreaks in poultry and wild birds and sometimes emerge into pandemic strains (Li et al., 2021; Ozawa and Kawaoka, 2013; Wille and Barr, 2022). HPAI viruses are of major importance as they have caused human infections, however, human-to-human transmission was negligible because of lack of virus adaptation. The recent 2.3.4.4b clade H5, a subclade of 2.3.4.4 viruses, was reassorted with N1 (H5N1) and caused devasting outbreaks throughout the year in wild birds and poultry (Gilbertson and Subbarao, 2023; Lewis et al., 2021; Xie et al., 2023). Intriguingly, it rapidly spread to several mammalian species and was successful in mammal-to-mammal transmission in farmed minks, New England seals and dairy cattle. Thus, the current H5N1 poses a potential pandemic threat if it gains the ability for human-to-human transmission. 2.3.4.4 clade H5, in addition to N1, also reassorted with multiple NAs (H5N2, H5N6, or H5N8) and led to previous outbreaks in poultry. Why did the combination of H5 and N1 of 2.3.4.4b clade reemerge as the dominant strain? The pairing of 2.3.4.4b clade H5 with the N1 is considered stochastic. However, our results here show that the evolution of glycan surface plasticity in 2.3.4.4b clade H5 HA unusually favored its pairing with the long-stalk N1 subtype and the evolution of the highly transmissible H5N1. Also, our results demonstrate a critical molecular principle driving the fitness of reassorted avian HA and NA subtypes. Why have avian H5N1 and H5N8 infections become prevalent in recent outbreaks? The 2.3.4.4 clade H5 HA, which emerged in 2014, is paired with multiple NA subtypes (N2, N6, and N8) (Gilbertson and Subbarao, 2023; Li et al., 2021; Yamaji et al., 2020). Only H5N6 and H5N8 stayed in circulation, and H5N8 emerged as the dominant strain until 2020. In 2020, H5N1 emerged from H5N8, which became dominant over all the 2.3.4.4b subtypes. Interestingly, N1 and N8 subtypes of NA, which paired with 2.3.4.4b clade H5 HA, consisted predominantly of longer stalk lengths, unlike N6, which showed short stalk length. Thus, 2.3.4.4b clade H5 was able to reassort with multiple NA subtypes, however, only NA with long stalk length conferred fitness advantages and further emergence of dominant circulating strain.

Why does the short stalk N1 subtype fail to confer fitness advantage for reassortment with 2.3.4.4b clade H5 HA? The HPAI H5 pairs with long and short-stalked N1, and the avian H5 that is reassorted with short stalk N1 dominated since the appearance of gs/GD lineage. However, the recent 2.3.4.4b clade H5 paired exclusively with long stalk NA and emerged as a dominant circulating strain, unlike the previous clades. Our observations indicate that short stalk N1 preferentially pairs with H5 with 8 or more glycosylation sites, whereas the long stalk N1 pairs with H5 having 7 or 6 glycosylation sites. This suggests an intricate association of H5 glycosylation motifs pattern with the stalk length of N1 for successful reassortment. The glycosylation site positions in H5 HA are highly conserved except at the 8^th^ glycosylation site position. The 8^th^ glycosylation site position in H5 varied across the time. Residue 103 in H5 HA marks the 8th glycosylation site early on, but this site was shifted later to residue 171, which is located near the sialic acid binding site. The currently circulating 2.3.4.4b clade consisted of 7 and 8 glycosylation site H5 variants, indicating its propensity to pair with long or short-stalk N1. However, only 2.3.4.4b clade H5 that paired with long-stalk N1 emerged as a dominant strain. This indicates that, although 2.3.4.4b H5 (with 8 glycosylation sites) reassorted with short stalk N1, this combination did not confer fitness advantage for H5N1 evolution. Perhaps the shift in the 8th glycosylation site position in 2.3.4.4b H5 might be the underlying fitness constraint for the successful pairing and emergence of H5N1 with short stalk N1. Also, the shift in 8^th^ glycosylation site in H5 preceded the deglycosylation event or evolution of 7 glycosylation sites variant of H5 that led to the emergence of 2.3.4.4b H5N1. Thus, a shift in the 8^th^ glycosylation site followed by deglycosylation and the emergence of the H5 variant of 7 glycosylation sites might underlie the 2.3.4.4b clade H5 pairing, preferably with the long stalk N1 subtype.

Is the pairing of avian HA and NA stochastic? The glycan surface plasticity of the H5 is critical for its pairing with NA subtypes of specific stalk lengths. The number of circulating H5 variants with diverse glycosylation site patterns enables pairing with multiple NA subtypes with short or long stalk lengths. Our results indicate that the glycosylation pattern of avian H5 and H9 show epistatic interaction with specific NA subtypes and their stalk lengths for reassortment and emergence as dominant strains in circulation. Also, we demonstrated that the glycosylation site patterns of avian HA and the stalk lengths of NA subtypes would act as molecular clues for predicting the evolution and emergence of novel avian influenza strains.

## METHODS

### Pipeline development and accessibility

We established a reproducible data analysis pipeline accessible via GitHub, designed to streamline data processing from initial FASTA file input to the production of figures presented in this study. The core of the pipeline is comprised of Jupyter notebooks, which are available within the repository ([Github repository](https://github.com/ranjanaanataraj/2.3.4.4b-H5N1-analysis)).

### Temporal analysis of IAV circulating strains across host types

To elucidate trends in circulating influenza strains and neuraminidase (NA) subtype prevalence, we gathered infection counts for each Influenza A Virus (IAV) subtype from the Global Initiative on Sharing All Influenza Data (GISAID) platform ([GISAID Link](https://platform.epicov.org/epi3/frontend#49c9ab)). Sequences were filtered based on host type, time period, and HA and NA types. We opted to visualize the normalized data instead of absolute numbers to avoid the potential bias arising from overrepresentation and facilitate comparisons across time periods. This normalization process was performed to create heatmaps using the Seaborn library (v0.13.1) in Python (v3.10.12).

### Construction of stalk length distributions

To construct the stalk length distributions of the NA subtypes, we first defined the domain boundaries of the consensus sequence for each NA sequence dataset. The stalk domain of the consensus sequence was delineated by aligning it with a domain-demarcated PR8 NA sequence using the Clustal Omega Server. Subsequently, the stalk length was computed by tallying the non-gap characters within the aligned region. To automate this analysis, custom Python scripts were developed, using the computational capabilities of numpy and Pandas libraries. The results of this analysis were visually depicted using kernel density estimate (KDE) plots in Seaborn.

### H3 Numbering

Amino acid positions were reported using the H3 numbering scheme. Alignment with the HA1 and HA2 chains of PDB 4HMG was executed using Clustal Omega, and the corresponding positions were documented.

### Glycosylation Site Analysis

The putative glycosylation site maps were constructed based on sequence data for avian influenza HA and NA proteins, which were fetched as FASTA files from the EpiFlu database within GISAID.org. These datasets were consolidated based on unique EPI_ISL_IDs to ensure that the HA and NA were from the same virus sample and to maintain the data integrity and accuracy. To identify putative glycosylation sites within the protein sequences, a custom Python script was developed, leveraging the functionality of Pandas and Regex libraries that facilitate large dataset analysis and pattern identification. This script systematically scanned the sequences for the specified pattern N-[A-Z]-[S/T], indicating potential glycosylation sites. The identified sites were then visualized using Seaborn’s kernel density estimate (KDE) plots, providing a comprehensive overview of the glycosylation landscape across the viral proteins. The glycosylation site positions were then annotated based on H3 sequence numbering. To calculate percentage conservation of glycosylation sites, the multiple sequence alignment datasets were generated and individual glycosylation sites were defined (as mentioned above) for scoring conservation of the glycosylation site. An intact glycosylation motif (N-X-T/S) was assigned a score of 1, and the sites mutated were scored as 0. Using this approach, the conservation of each glycosylation site was calculated and represented in heat maps (Seaborn library (v0.13.1)).

To track the percent conservation of H5 glycosylation sites in avian species, we used influenza virus sequence data from GISAID after filtering for wild aquatic birds (geese, gulls, mallad ducks, and swan) and domestic birds (chicken) (Olsen et al., 2006; Xie et al., 2023).

### Multiparametric Bubble Plots of avian influenza HA and NA Pairing

We employed the scatter plot function of Plotly Express (v5.15.0) to generate the bubble plots showcased in Figure 6. Each point on the plot represents four molecular features of the H5/H9 and NA subtypes pairing. Specifically, the number of glycosylation sites of HA was depicted as the x-axis parameter, and the NA subtype paired with the particular HA number of glycosylation sites of H5/H9 HA was encoded on the y-axis.

The frequency of these combinations was visualized using the size argument of the scatter function, providing insight into the prevalence of specific avian influenza HA and NA pairings. Additionally, we encoded the stalk length information of specific NA subtypes using the color parameter. The HxNy sequence data utilized in this analysis was obtained from GISAID as FASTA files. To determine the putative glycosylation coverage of the H5/H9 HA sequences, we followed the methodology outlined in the glycosylation site analysis section of the methods. Similarly, for each NA type, the stalk length was calculated as detailed previously.

### Structural Analysis

To analyze the glycosylation site positions within each H5 dataset, we first identified these glycosylation motifs within the consensus sequence. Subsequently, we mapped these identified glycosylation sites onto the experimentally solved structure of trimeric avian H5 HA bound with sialic acid in the pocket (PDB: 3UBE). This annotation aided in understanding the spatial distribution of glycosylation sites and their potential functional implications within the H5 protein structure. For both structural analysis and annotations, we utilized UCSF Chimera (v1.17.3). Each visual representation of the H5 HA structure was annotated to highlight key domains - the Receptor Binding Domain (RBD) and the Vestigial Esterase (VE) domain. The domain boundaries were defined as follows: RBD – residues 136-254, and VE – residues 59-135 and 256-262. To provide a clear depiction of the H5 HA with glycosylation sites, we represented the position of the beginning of the glycosylation motif (N residue of NxS/T) as spheres within the structure.

## ACKNOWLEDGEMENTS

We thank S.K. lab members for providing critical comments. We thank Keerthana Balamurugan from IISc for helping initial GISAID sequence analysis. We sincerely thank Richard J. Webby from St. Jude Children’s Research Hospital for his critical suggestions during the preparation of this manuscript. This work was supported by funding from the Department of Biotechnology (DBT) (BT/PR45145/COT/142/24/2022), Indian Council of Medical Research (ICMR) (EMDR/SG/9/2023-2019) (2021-14148/CMB/ADHOC-BMS), Science and Engineering Research Board (SERB-DST) (EEQ/2021/000274), Scheme for Transformational and Advanced Research in Sciences (STARS)-MoE grant (MoE-STARS/STARS-2/2023-0464) and the Indian Institute of Science (IISc) research support.

## AUTHOR CONTRIBUTIONS

S.K. conceptualized the study; S.K., R.N., and A.C. analysed and investigated the study; R.N. developed methods to analyse large sequence data sets; S.K., R.N., and A.C. wrote the manuscript and edited the manuscript; S.K. supervised the study; S.K. provided the guidance and brought the funding.

## SUPPLEMENTARY FIGURE LEGENDS

**Figure S1: The analysis of NA subtypes in circulation and their association with avian H5 HA: A.** Pie chart illustrating the prevalence of IAV neuraminidase subtypes deposited in the GISAID server from 1990-2023. Color-coded sectors represent distinct IAV subtypes, with annotations indicating the percentage of sequences. **B.** Pie chart illustrating the distribution of avian Influenza A Virus (IAV) subtypes deposited in the GISAID server from 1990 to 2023. The sectors are color-coded according to the IAV subtype and annotated by the percentage of sequences deposited. **C.** Decade-wise trends in circulating IAV strains in avian hosts represented as heatmaps. The horizontal axis displays the hemagglutinin (HA) subtypes (H1-H11), while the vertical axis shows the NA subtypes (N1-N9). Each cell in the heatmap denotes a specific IAV subtype, annotated with the frequency of its sequences represented in GISAID and color-coded accordingly. Heatmaps are provided for four distinct time periods: 1990-1999, 2000-2009, 2010-2019, and 2020-2023. **D, E.** Decade-wise heatmaps similar to Panel-C constructed for swine and human host types, respectively.

**Figure S2: Long stalk N1 and N8 preference for pairing with 2.3.4.4b clade H5 HA. A.** KDE plot representing the distribution of N1 stalk lengths in clade 2.3.4.4b H5N1 infections in bobcats, foxes, dolphins, and penguins. **B.** Consensus protein sequence alignment of N1 stalk regions of avian, human, and cow infected with 2.3.4.4b clade H5N1 viruses. **C**. Stalk length distribution of all avian N2 subtype sequences deposited in GISAID from 1990 to 2023, presented as kernel density (KDE) plots. Each time period is depicted as a separate curve. **D.** Consensus protein sequence alignment of the two long stalk variants of the N2 subtype in avians. The putative glycosylation motif (Nx[S/T]) differentiating these two variants is highlighted in yellow.

**Figure S3: Amino acid modifications in avian N1 head domain from 2010-19 and 2020-2023**. **A**. AlphaFold2 predicted structures of NA head domain consensus sequences from 2010-19 and 2020-23. **B**. A close-up representation of the active site pocket of predicted N1 head domain from H5N1 sequences 2010-19. The residues essential for sialic acid binding are highlighted in cyan and their side chains are represented in sticks. The sialic acid moiety is shown in yellow and colored by heteroatoms. **C**. Active site residues of predicted N1 head domain from 2020 shown similar to panel B. **D**. Table showing the residue positions mutated in N1 head domain from 2010-19 and 2020-23. **E**. The residues from panel D (in red spheres) mapped on to the crystal structure of N1 head domain (PDB ID: 2HTY) and the C_α_-C_β_ atoms of the active site residues essential for sialic acid binding are highlighted in cyan. **F**. Closeup view of the active site pocket from Panel E.

**Figure S4: The pairing of long stalk N1 confers better fitness with 2.3.4.4b clade H5 compared to the long stalk N8**. **A**. The pie charts illustrate the pairing of avian H5Nx strains observed from 2010 to 2019. On the left, the distribution for clade 2.3.4.4b H5 is shown, while on the right, the cumulative distribution over all H5 clades in circulation is depicted. The sectors are color-coded by the NA subtype and annotated by the percentage of sequences they constitute. **B.** The KDE plot depicts the distribution of stalk lengths of the NA subtypes - N1, N6, and N8 that were found to pair with H5 strains, as illustrated in Panel-A. The color scheme of the density profile follows that of Panel-A. **C**. The frequency distribution of H5Nx sequences is plotted for the time period spanning from 2020 to 2023, similar to Panel-A. **D**. The stalk length distribution of NA subtypes that are paired with H5 is constructed similarly to Panel B. **E**. The pie charts illustrate the clade composition of avian H5N1, H5N6, and H5N8 sequences from 2014-2023. The sectors are color-coded by H5 HA clades. The size of the pie chart reflects the abundance of that strain in circulation.

**Figure S5: Annotating variations in 2.3.4.4b clade H5 HA that paired with either short stalk or long stalk N1 subtype. A.** Amino acid substitutions in H5 HA associated with N1 stalk length variation. Illustration of 43 amino acid substitutions in H5, distinguished by its pairing with short-stalked N1 (left) and the corresponding residues when paired with long-stalked N1. The amino acid positions are indicated in H3 numbering. **B.** The table representing sequence coverage for avian species used for the analysis in Figure-2B.

**Figure S6: The evolution of glycosylation sites in domestic and wild aquatic birds**. Heatmap depicting the percent conservation of glycosylation sites of the H5 HA obtained from a specific time frame as indicated in the plot for domestic (**A**) and wild aquatic birds (**B**). The horizontal axis shows the years sampled, and the vertical axis indicates glycan positions of H5 in H3 numbering. We analysed 4075 sequences of wild birds (swans, geese and mallard ducks) and 3138 sequences of domestic birds (chicken and poultry ducks) from GISAID.

**Figure S7: The requirement of seven glycosylation site variant of 2.3.4.4b clade H5 to pair with long stalk N1 and N8 subtypes**. **A.** KDE plot illustrates the number of putative glycosylation sites in clade 2.3.4.4b H5 paired with N6 and N8 subtypes during 2010-2019. **B.** KDE plot constructed using 2.3.4.4b clade H5N1 and H5N8 sequences from 2020 to 2023. **C**. Impact of 8th glycosylation site loss in avian H5. The H5 HA trimer, visualized using the structure (PDB: 3UBE), with one monomer color-coded by domains – RBD in green, VE domain in yellow, F1 region in blue, and HA2 stalk in orange. The remaining monomers are depicted in grey. Zoomed-in insets reveal atomic contacts at position 105, showcasing potential interactions with an intact glycosylation site (top inset) and altered atomic contacts when the T105 is mutated to E (bottom inset).

