## Supplementary figures for "Avian influenza virus neuraminidase stalk length and haemagglutinin glycosylation patterns reveal molecularly directed reassortment promoting the emergence of highly pathogenic clade 2.3.4.4b A (H5N1) viruses"

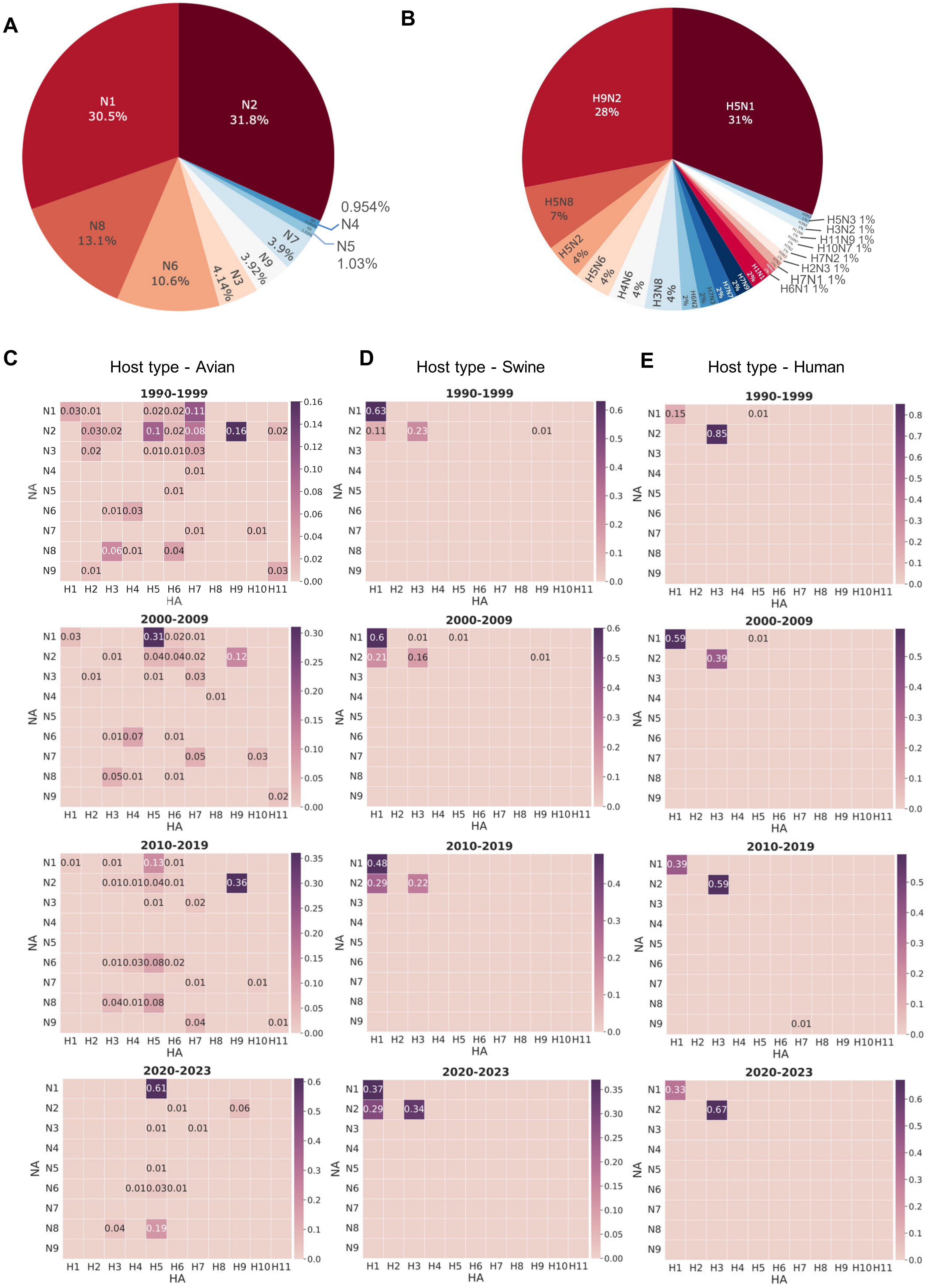
A
N1
30.5%
N2
31.8%
N8
13.1%
N6
10.6%
N3
4.14%
N9
3.92%
N7
3.9%
0.954%
N4
N5
1.03%
B
H9N2
28%
H5N1
31%
H5N8
7%
H5N2
4%
H5N6
4%
H4N6
4%
H3N8
4%
H6N2
2%
H7N3
2%
H7N7
2%
H7N9
2%
H1N1
2%
H5N3 1%
H3N2 1%
H11N9 1%
H10N7 1%
H7N2 1%
H2N3 1%
H7N1 1%
H6N1 1%
C
Host type - Avian
D
Host type - Swine
E
Host type - Human
1990-1999
2000-2009
2010-2019
2020-2023
NA
HA
N1
N2
N3
N4
N5
N6
N7
N8
N9
H1 H2 H3 H4 H5 H6 H7 H8 H9 H10 H11

**Figure S1**

**A**

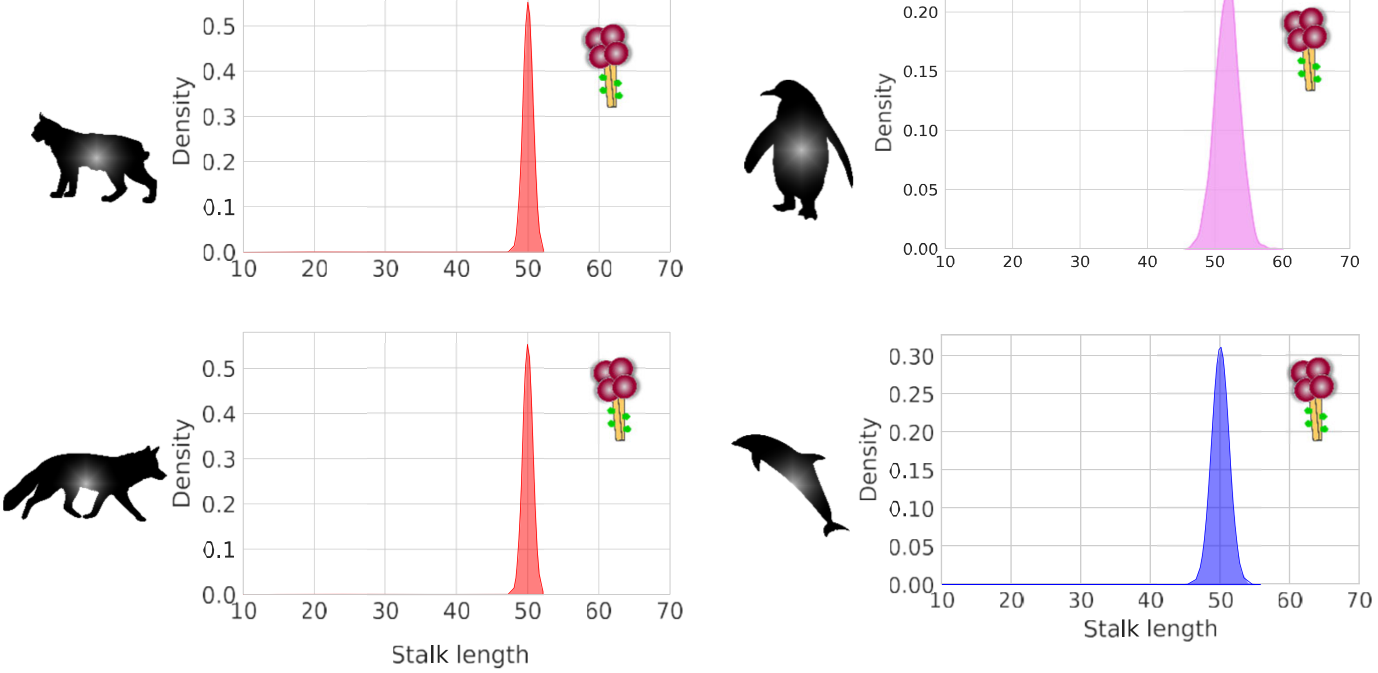


**B**

```
Avian_H5N1_N1_short_stalk          ISIWVSHSIQTGNQHQTEPIRNTNFLTEN--------------------AVAS 33
Cow_2.3.4.4b_H5N1_N1_stalk         ISIWVSHSIQTGNQYQPEPCNQSIITYENNTWVNQTYINISSTNFLAEQAVTS 53
Human_2.3.4.4b_H5N1_N1_stalk       ISIWVSHSIQTGNQYQPEPCNQSIITYENNTWVNQTYVNISNTNFLAEQAVTS 53
Avian_H5N1_N1_long_stalk           ISIWVSHSIQTGNQYQPEPCNQSIITYENNTWVNQTYVNISNTNFLAEQAVTS 53
```

**C**

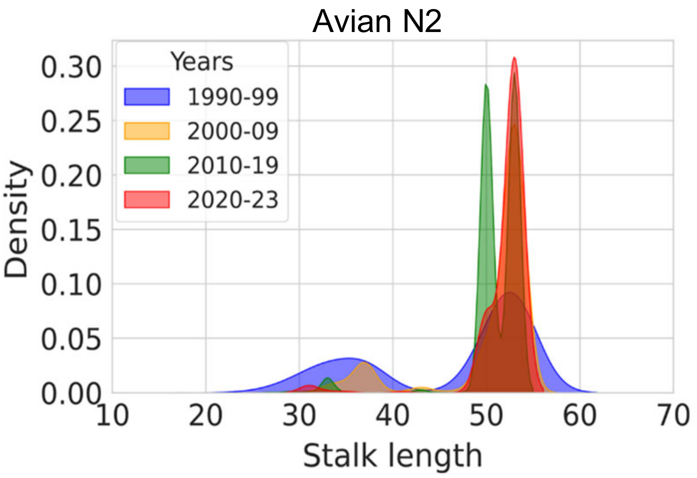


**D**

```
Avian_N2_53_stalk      ATTVTLHFKQNECSIPxNNQVVPCEPIIIERNITEIVYLNNTTIEKEICPKVx      53
Avian_N2_50_stalk      TTTMTLHFGQKECSNPSNNQVMPCEPIIIERNT---VHLNSTTIEREICPKVA      50
```

**Figure S2**

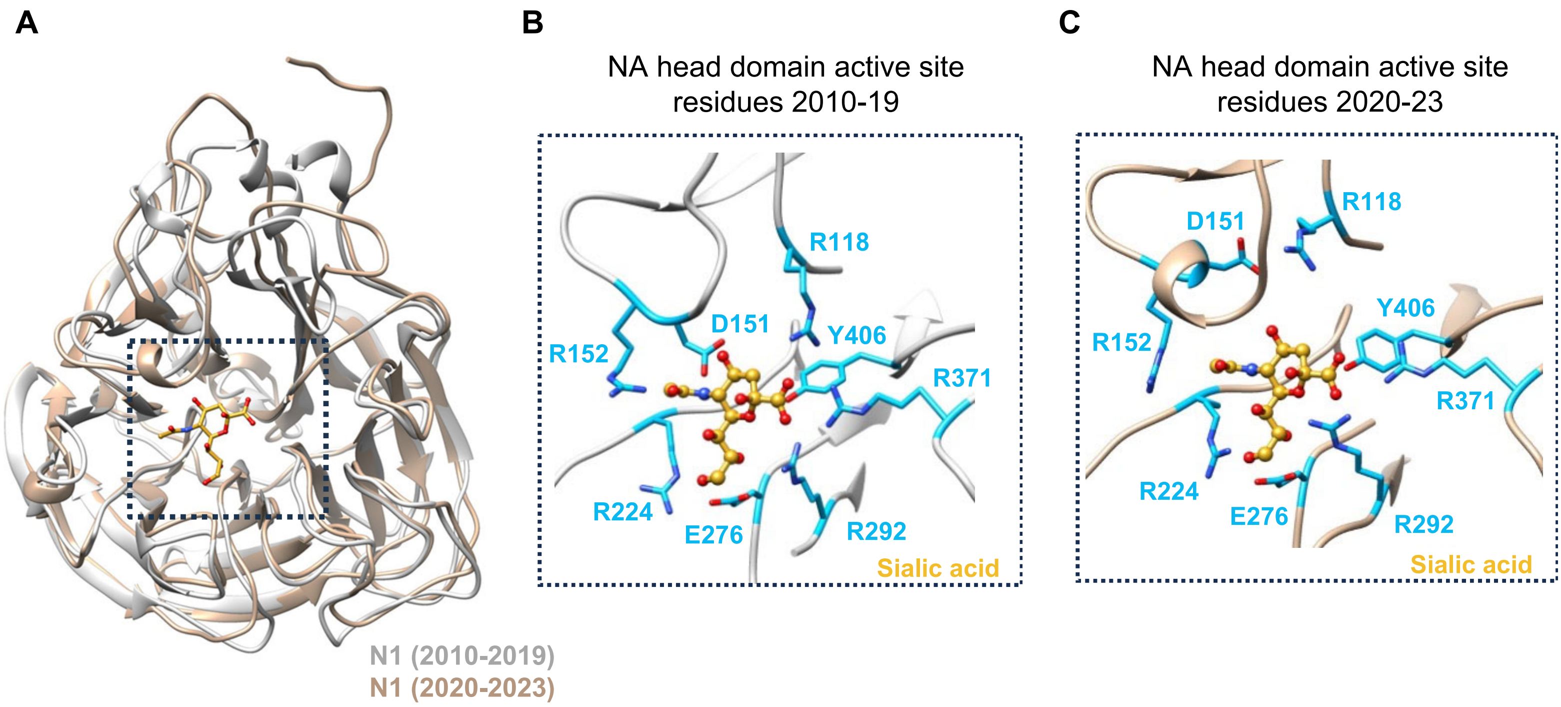


D

| | | |
|---|---|---|
| R13S | T106I | P258S |
| V17I | M176I | I264V |
| H18Y | L187M | N284S |
| S23G | E205D | G299E |
| E46D | T207M | E303S |
| S72P | G254S | A312E |
| H73Y | V256M | I335M |
| M81V | S257P | S351N |

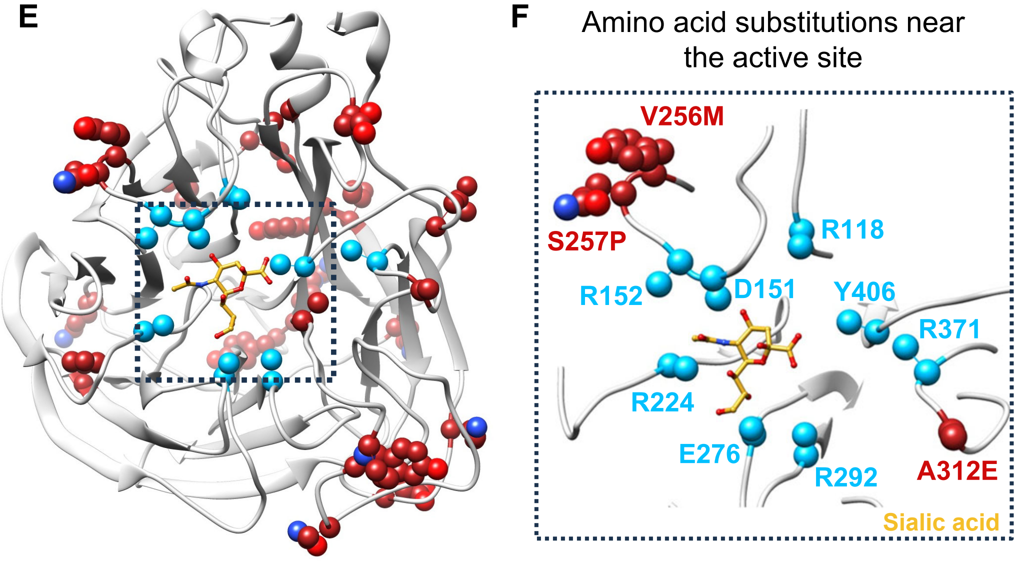


**Figure S3**

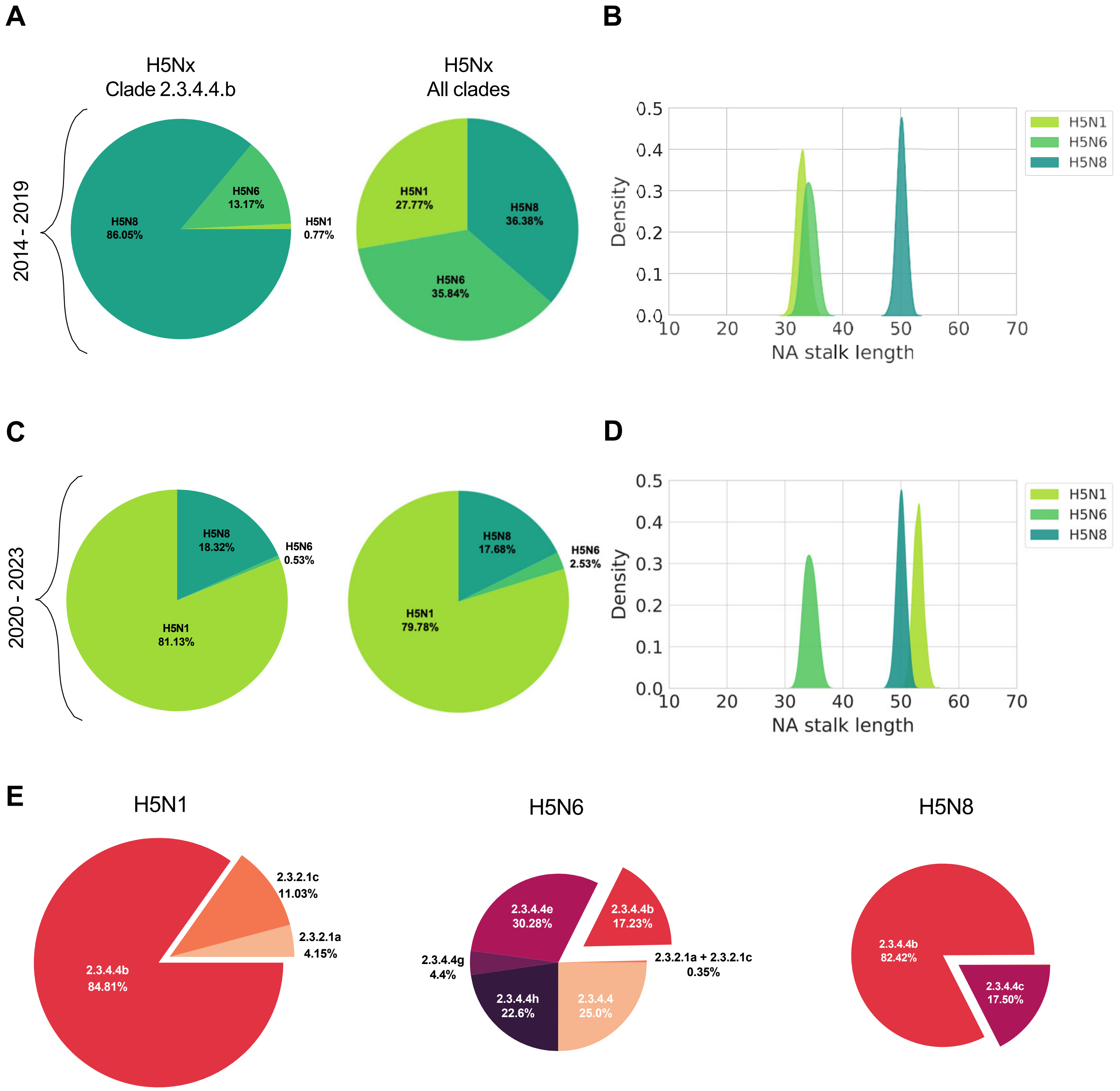
A
H5Nx
Clade 2.3.4.4.b
H5Nx
All clades
2014 - 2019
H5N8
86.05%
H5N6
13.17%
H5N1
0.77%
H5N1
27.77%
H5N8
36.38%
H5N6
35.84%
B
Density
NA stalk length
H5N1
H5N6
H5N8
C
2020 - 2023
H5N8
18.32%
H5N6
0.53%
H5N1
81.13%
H5N8
17.68%
H5N6
2.53%
H5N1
79.78%
D
Density
NA stalk length
H5N1
H5N6
H5N8
E
H5N1
2.3.2.1c
11.03%
2.3.2.1a
4.15%
2.3.4.4b
84.81%
H5N6
2.3.4.4e
30.28%
2.3.4.4b
17.23%
2.3.4.4g
4.4%
2.3.2.1a + 2.3.2.1c
0.35%
2.3.4.4h
22.6%
2.3.4.4
25.0%
H5N8
2.3.4.4b
82.42%
2.3.4.4c
17.50%

**Figure S4**

**A**

| | | | |
|---|---|---|---|
| F2L | S128P | E188A | E271G |
| H12Q | D129N | A189E | N276H |
| N81R | A132T | R193N | R280K |
| K90R | N144A | Q196K | I285V |
| N101S | S145P | I204V | Q325L |
| F102L | N159D | K222Q | K455R |
| R122L | K166I | I223V | |
| D125S | S173A | S227R | |
| | Q186R | I230M | |
| | D187N | N240D | |

**B**

| Avian species | 2.3.2.1a 2.3.2.1c | 2.3.4.4b |
|---|---|---|
| **Wild** | 390 | 6085 |
| **Domestic** | 729 | 1929 |
| **Duck** | 818 | 250 |

**Figure S5**

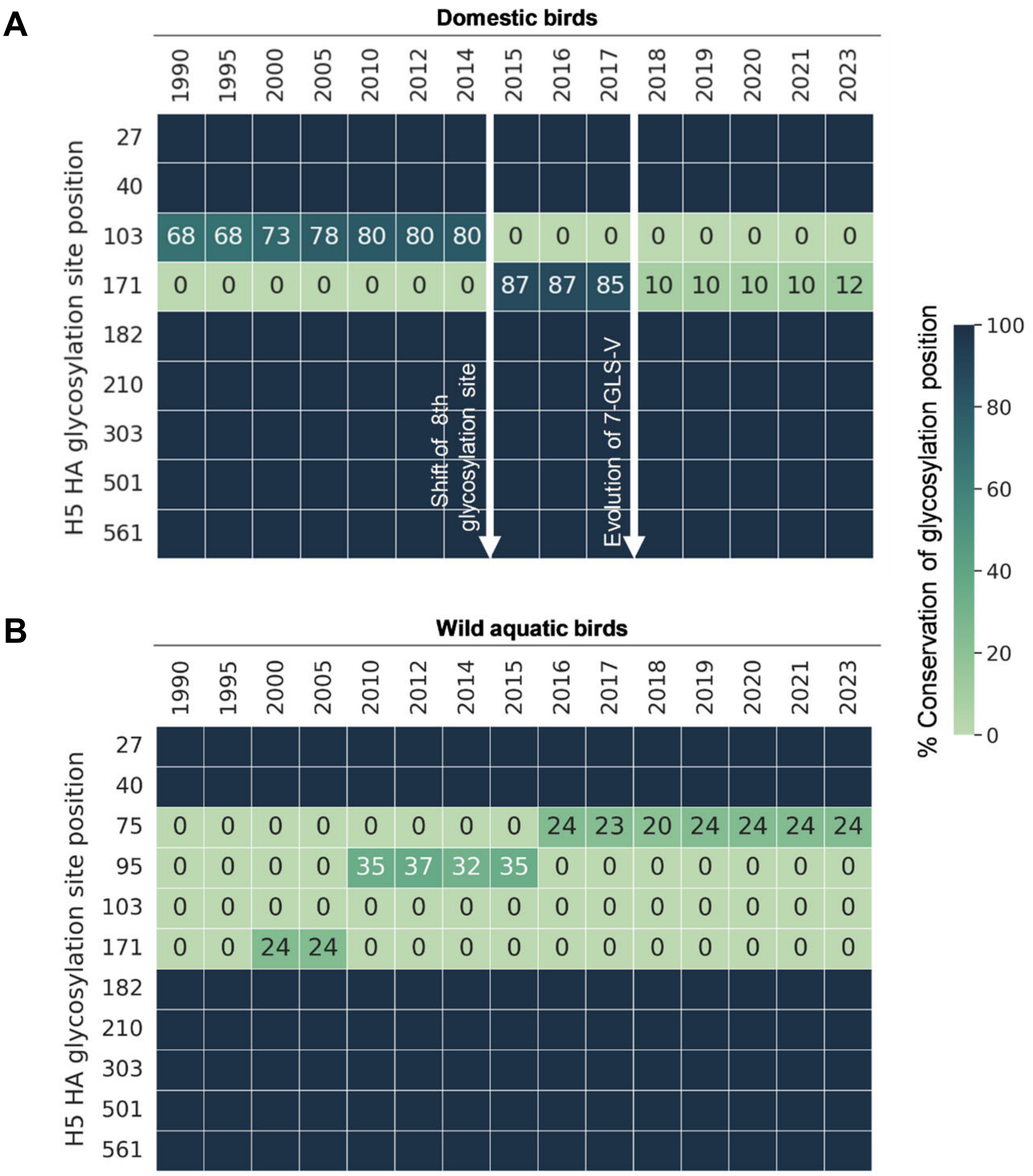
A
Domestic birds
1990 1995 2000 2005 2010 2012 2014 2015 2016 2017 2018 2019 2020 2021 2023
H5 HA glycosylation site position
27
40
103: 68 68 73 78 80 80 80 0 0 0 0 0 0 0 0
171: 0 0 0 0 0 0 0 87 87 85 10 10 10 10 12
182
210
303
501
561
Shift of 8th glycosylation site
Evolution of 7-GLS-V
B
Wild aquatic birds
1990 1995 2000 2005 2010 2012 2014 2015 2016 2017 2018 2019 2020 2021 2023
H5 HA glycosylation site position
27
40
75: 0 0 0 0 0 0 0 0 24 23 20 24 24 24 24
95: 0 0 0 0 35 37 32 35 0 0 0 0 0 0 0
103: 0 0 0 0 0 0 0 0 0 0 0 0 0 0 0
171: 0 0 24 24 0 0 0 0 0 0 0 0 0 0 0
182
210
303
501
561
% Conservation of glycosylation position
100
80
60
40
20
0

**Figure S6**

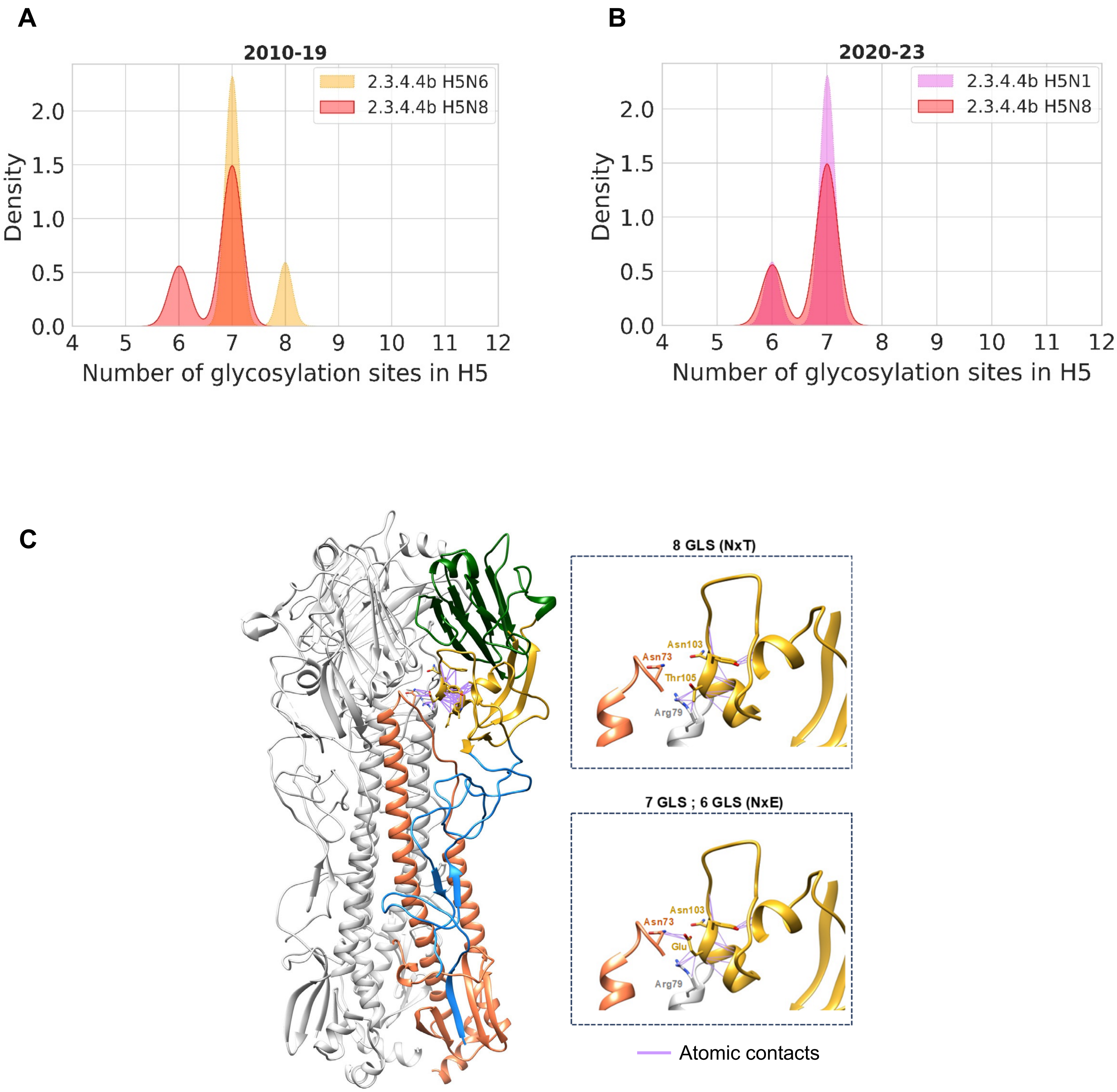
A
2010-19
2.3.4.4b H5N6
2.3.4.4b H5N8
Density
0.0
0.5
1.0
1.5
2.0
4 5 6 7 8 9 10 11 12
Number of glycosylation sites in H5
B
2020-23
2.3.4.4b H5N1
2.3.4.4b H5N8
Density
0.0
0.5
1.0
1.5
2.0
4 5 6 7 8 9 10 11 12
Number of glycosylation sites in H5
C
8 GLS (NxT)
Asn103
Asn73
Thr105
Arg79
7 GLS ; 6 GLS (NxE)
Asn103
Asn73
Glu
Arg79
Atomic contacts

Figure S7
